# Unique Evolution of Antiviral Tetherin in Bats

**DOI:** 10.1101/2020.04.08.031203

**Authors:** Joshua A. Hayward, Mary Tachedjian, Adam Johnson, Aaron T. Irving, Tamsin B. Gordon, Jie Cui, Alexis Nicolas, Ina Smith, Victoria Boyd, Glenn A. Marsh, Michelle L. Baker, Lin-Fa Wang, Gilda Tachedjian

## Abstract

Bats are recognised as important reservoirs of viruses deadly to other mammals, including humans. These infections are typically non-pathogenic in bats raising questions about host response differences that might exist between bats and other mammals. Tetherin is a restriction factor which inhibits the release of a diverse range of viruses from host cells, including retroviruses, coronaviruses, filoviruses, and paramyxoviruses, some of which are deadly to humans and transmitted by bats. Here we characterise the tetherin genes from 27 species of bats, revealing that they have evolved under strong selective pressure, and that fruit bats and vesper bats express unique structural variants of the tetherin protein. Tetherin was widely and variably expressed across fruit bat tissue-types and upregulated in spleen tissue when stimulated with Toll-like receptor agonists. The expression of two computationally predicted splice isoforms of fruit bat tetherin was verified. We identified an additional third unique splice isoform which includes a C-terminal region that is not homologous to known mammalian tetherin variants but was functionally capable of restricting the release of filoviral virus-like particles. We also report that vesper bats possess and express at least five tetherin genes, including structural variants, a greater number than any other mammal reported to date. These findings support the hypothesis of differential antiviral gene evolution in bats relative to other mammals.

## Introduction

Bats are reservoirs of viruses that are highly pathogenic to other mammals, including humans. Viruses such as Hendra, Nipah, Ebola, Marburg, severe acute respiratory syndrome coronaviruses (SARS-CoV-1 and likely also SARS-CoV-2), and the Sosuga virus have crossed species barriers from bats into humans (1–5). Laboratory studies demonstrate that specific bat species can be infected with Ebola, Marburg, SARS-CoV-1, Hendra, and Nipah viruses without showing clinical signs of disease (6–10). Recently, this has led to efforts to elucidate if there are differences observed between the antiviral strategies of bats and other mammals (11–19). These studies have identified that genes related to immunity in bats are under a significant degree of positive selection, in addition to differences in the copy number and diversity of innate immune genes of bats relative to other mammals (12-15, 17, 20).

Tetherin (BST-2, CD317) is a mammalian restriction factor that inhibits the release of diverse enveloped viral particles from the cells in which they are produced (21–23). These viruses include retroviruses, coronaviruses, filoviruses, and paramyxoviruses, some of which are transmissible by bats (24–27). Tetherin is a membrane-associated glycoprotein and its activity is determined by its structure, rather than its primary amino acid sequence (28, 29). In addition, tetherin possesses a secondary role in immune signalling events by triggering the NF-κB signalling pathway, leading to stimulation of the antiviral interferon response (30–33).

In humans, tetherin is expressed across most cell types, including its BST-2 namesake bone marrow stromal cells, and its expression is upregulated by stimulation with type I interferons (34–36). Tetherin is a dimeric dual-anchor type II membrane protein that contains one protein anchor, a transmembrane domain near its N-terminus, an extracellular coiled-coil domain, and a glycophosphatidylinositol (GPI) lipid anchor, which is attached to its C-terminus as a post-translational modification (37–39). Tetherin contains a number of conserved cytosine and asparagine motifs within the extracellular domain, with respective roles in dimerisation and glycosylation, and a dual-tyrosine motif (Y·x·Y) in its cytoplasmic region, which has a role in viral particle endocytosis and immune signalling cascades (31, 40). Tetherin is located in lipid rafts at the plasma membrane where many viral particles bud during acquisition of their host membrane-derived viral envelopes (39). During the viral budding process one anchor remains embedded in the nascent viral envelope while the other remains attached to the plasma membrane, tethering the virion to the cell and preventing its release into the extracellular environment (28, 38).

Tetherin is common to all mammals, and orthologs which share structural and functional, but not sequence similarity, exist in other vertebrates (29). Most mammals carry only a single tetherin gene; however gene duplication has been observed in sheep, cattle, opossums and wallabies (29, 41). Furthermore, human tetherin is expressed in two alternative isoforms (30), the long (l-tetherin) and short (s-tetherin) isoforms, which differ through the truncation of 12 amino acid (AA) residues at the N-terminus of l-tetherin. Computational analysis of the genomes of megabats from the *Pteropus* genus predict two additional isoforms, X1 and X2, with an internal, rather than N-terminal difference in amino acid sequences, although whether these isoforms are expressed is unknown (12). The predicted isoform X1 is homologous to human l-tetherin, and the X2 isoform is a splice variant which contains a 7 AA exclusion within the extracellular coiled-coil domain relative to isoform X1. A recent analysis of tetherin in the fruit bats *Hypsignathus monstrosus* and *Epomops buettikoferi* revealed that these species expressed a homolog of *Pteropus* isoform X1 and that it is a functional restriction factor capable of inhibiting the release of the bat-hosted Nipah virus, as well as virus-like particles (VLPs) and GP-virosomes produced from the Ebola virus and the human immunodeficiency virus (23, 42).

We hypothesised that given the role of bats as hosts of pathogenic enveloped viruses, and the differences observed in other antiviral genes of their innate immune system (14), that diversity might also exist in the tetherin genes of bats. Here we report that fruit bats possess a unique structural isoform which has differential activity against VLPs, restricting filoviral but not retroviral VLPs, in addition to the two computational predictions, tetherin isoforms X1 and X2, which were generated through the automated NCBI annotation pipeline from the published *P. alecto* genome [NCBI: PRJNA171993] (12). Adding to a previous report describing the presence and expression of three tetherin paralogs in the microbat *Myotis daubentonii,* and four tetherin genes in the genome of *M. lucifugus* (suborder Yangochiroptera) (18) we have found that microbats of the genus *Myotis* (e.g. *M. macropus*) possess at least five tetherin genes, two of which contain large structural differences in the extracellular coiled-coil domain. An analysis of the tetherin genes of 27 species of bats (including *P. alecto*) revealed that bat tetherin has been subjected to strong positive selection, indicating that these flying mammals have experienced evolutionary pressure on this antiviral gene. Collectively our findings indicate that bats have undergone tetherin gene expansion and diversification relative to other mammals, supporting the hypothesis of differential antiviral gene evolution in bats.

## Results

### Bats possess structural homologs of human tetherin

To advance our understanding of bat tetherin homologs, we mined all available publicly accessible sequence read archives representing 26 species of bats through a BLASTn analysis using the *P. alecto* tetherin isoform A as the query sequence (Table 1). The species analysed represent 8 of the 20 families within the order Chiroptera: three families (Hipposideridae, Pteropodidae, and Rhinolophidae) from within the suborder Yinpterochiroptera, and five families (Emballonuridae, Miniopteridae, Molossidae, Vespertilionidae, and Phyllostomidae) from within the suborder Yangochiroptera. This analysis revealed mRNA sequences representing homologs of human tetherin from 26 bat species, in addition to the homolog obtained from *P. alecto* cDNA, enabling the generation of consensus bat tetherin structural domains and amino acid sequence (Figure 1A & 1C) from a multiple sequence alignment (SI Data 1). The primary sequence lengths range from 177 to 221 AA, compared to 180 AA for human l-tetherin, and of these only 23 residues were conserved in 100% of the bat tetherin sequences analysed (Figure 1B). The assembled tetherin sequence for all bats was a homolog of human l-tetherin, except for the Black-bearded Tomb bat (*Taphozous melanopogon*), which was predicted to possess an equivalent of the human s-tetherin. The predicted *T. melanopogon* tetherin nucleotide sequence contains two key mutations upstream of its short isoform start site, an ATG to CTG change in the long isoform start codon, and a TAG stop codon in between the two start codon sites, either of which would be sufficient to prevent the production of the long isoform of tetherin. No nucleotide conflicts exist between the 17 reads mapping to this region of the *T. melanopogon* tetherin sequence, indicating that these mutations are unlikely to represent sequencing or consensus alignment errors.

**Figure 1.**
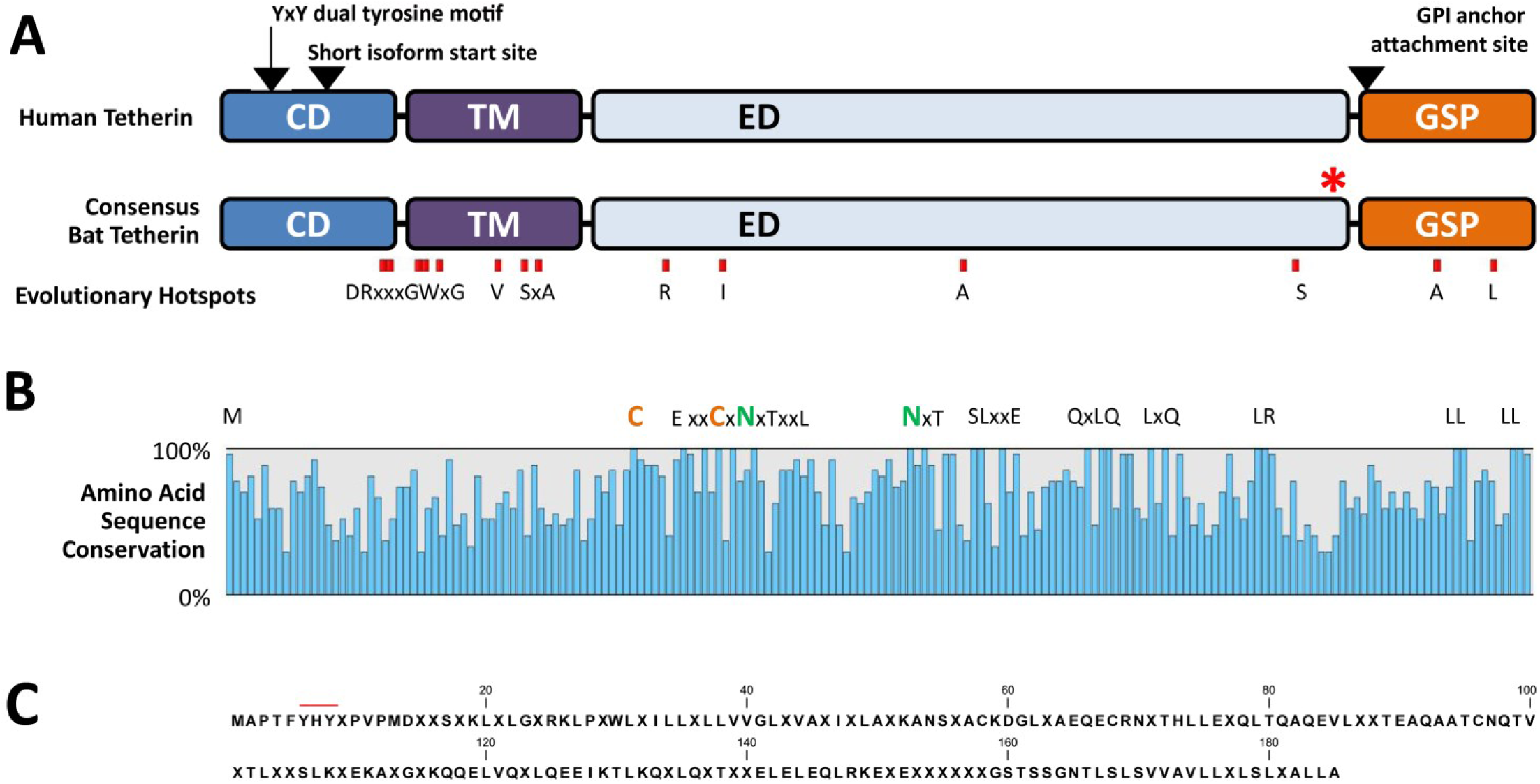
Bat Tetherin structure and sequence diversity. **A.** The consensus bat tetherin amino acid sequence structures and motifs generated through the multiple sequence alignment of tetherin from 27 bat species (SI Data 1) and compared against human tetherin. Amino acid residues under strong positive selection are denoted by red boxes. The YxY dual tyrosine motif, the alternative start site for the short isoform of tetherin, and the GPI anchor attachment site positions are indicated by arrows. The red asterisk (*) indicates the location of the inserted molecular tags in the *P. alecto* and *M. macropus* tetherin expression constructs. CD, cytoplasmic domain; TM, transmembrane domain; ED, extracellular domain; GPI, glycophosphatidylinositol; GSP, GPI signal peptide. **B.** Sequence diversity among bat tetherin proteins is indicated by the percentage of amino acid sequence conservation at each site of the consensus bat tetherin. Amino acids conserved in all 27 sequences are represented by their letters. Amino acids represented by X are variable and are included to indicate the sequence distance between closely positioned conserved residues. **C.** The consensus bat tetherin sequence. Amino acid residues represented in > 50% of bat tetherin sequences are indicated with their letter. Positions at which no amino acid residue is represented in > 50% of bat tetherin sequences are indicated with ‘X’. The red line indicates the position of the dual tyrosine motif.

**Table 1.**
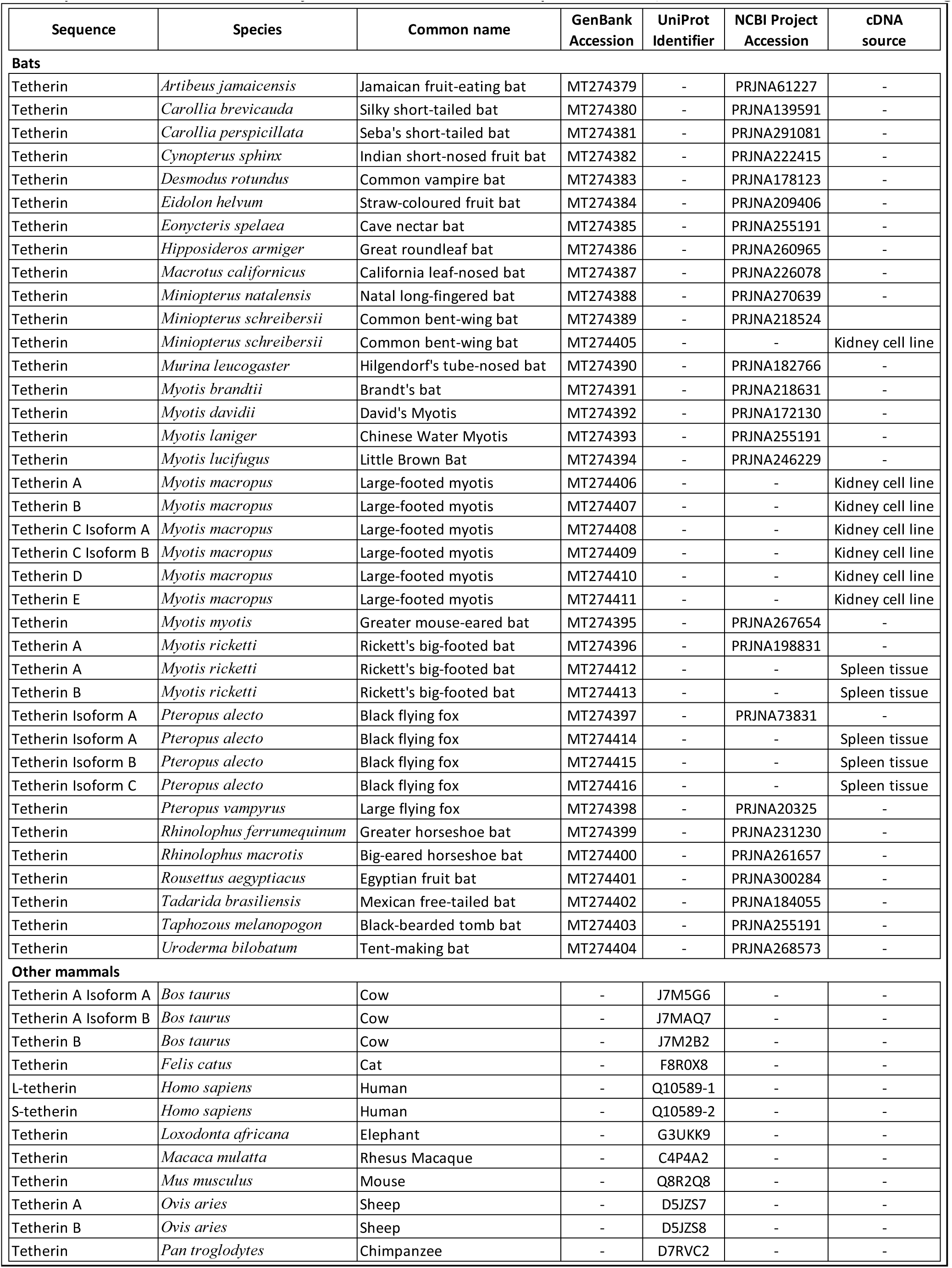
Sequence accessions with BioProject and cDNA sources for the prediction and/or confirmation of tetherin homologues

| Sequence | Species | Common name | GenBank Accession | UniProt Identifier | NCBI Project Accession | cDNA source |
| --- | --- | --- | --- | --- | --- | --- |
| **Bats** |  |  |  |  |  |  |
| Tetherin | *Artibeus jamaicensis* | Jamaican fruit-eating bat | MT274379 | - | PRJNA61227 | - |
| Tetherin | *Carollia brevicauda* | Silky short-tailed bat | MT274380 | - | PRJNA139591 | - |
| Tetherin | *Carollia perspicillata* | Seba's short-tailed bat | MT274381 | - | PRJNA291081 | - |
| Tetherin | *Cynopterus sphinx* | Indian short-nosed fruit bat | MT274382 | - | PRJNA222415 | - |
| Tetherin | *Desmodus rotundus* | Common vampire bat | MT274383 | - | PRJNA178123 | - |
| Tetherin | *Eidolon helvum* | Straw-coloured fruit bat | MT274384 | - | PRJNA209406 | - |
| Tetherin | *Eonycteris spelaea* | Cave nectar bat | MT274385 | - | PRJNA255191 | - |
| Tetherin | *Hipposideros armiger* | Great roundleaf bat | MT274386 | - | PRJNA260965 | - |
| Tetherin | *Macrotus californicus* | California leaf-nosed bat | MT274387 | - | PRJNA226078 | - |
| Tetherin | *Miniopterus natalensis* | Natal long-fingered bat | MT274388 | - | PRJNA270639 | - |
| Tetherin | *Miniopterus schreibersii* | Common bent-wing bat | MT274389 | - | PRJNA218524 | - |
| Tetherin | *Miniopterus schreibersii* | Common bent-wing bat | MT274405 | - | - | Kidney cell line |
| Tetherin | *Murina leucogaster* | Hilgendorf's tube-nosed bat | MT274390 | - | PRJNA182766 | - |
| Tetherin | *Myotis brandtii* | Brandt's bat | MT274391 | - | PRJNA218631 | - |
| Tetherin | *Myotis davidii* | David's Myotis | MT274392 | - | PRJNA172130 | - |
| Tetherin | *Myotis laniger* | Chinese Water Myotis | MT274393 | - | PRJNA255191 | - |
| Tetherin | *Myotis lucifugus* | Little Brown Bat | MT274394 | - | PRJNA246229 | - |
| Tetherin A | *Myotis macropus* | Large-footed myotis | MT274406 | - | - | Kidney cell line |
| Tetherin B | *Myotis macropus* | Large-footed myotis | MT274407 | - | - | Kidney cell line |
| Tetherin C Isoform A | *Myotis macropus* | Large-footed myotis | MT274408 | - | - | Kidney cell line |
| Tetherin C Isoform B | *Myotis macropus* | Large-footed myotis | MT274409 | - | - | Kidney cell line |
| Tetherin D | *Myotis macropus* | Large-footed myotis | MT274410 | - | - | Kidney cell line |
| Tetherin E | *Myotis macropus* | Large-footed myotis | MT274411 | - | - | Kidney cell line |
| Tetherin | *Myotis myotis* | Greater mouse-eared bat | MT274395 | - | PRJNA267654 | - |
| Tetherin A | *Myotis ricketti* | Rickett's big-footed bat | MT274396 | - | PRJNA198831 | - |
| Tetherin A | *Myotis ricketti* | Rickett's big-footed bat | MT274412 | - | - | Spleen tissue |
| Tetherin B | *Myotis ricketti* | Rickett's big-footed bat | MT274413 | - | - | Spleen tissue |
| Tetherin Isoform A | *Pteropus alecto* | Black flying fox | MT274397 | - | PRJNA73831 | - |
| Tetherin Isoform A | *Pteropus alecto* | Black flying fox | MT274414 | - | - | Spleen tissue |
| Tetherin Isoform B | *Pteropus alecto* | Black flying fox | MT274415 | - | - | Spleen tissue |
| Tetherin Isoform C | *Pteropus alecto* | Black flying fox | MT274416 | - | - | Spleen tissue |
| Tetherin | *Pteropus vampyrus* | Large flying fox | MT274398 | - | PRJNA20325 | - |
| Tetherin | *Rhinolophus ferrumequinum* | Greater horseshoe bat | MT274399 | - | PRJNA231230 | - |
| Tetherin | *Rhinolophus macrotis* | Big-eared horseshoe bat | MT274400 | - | PRJNA261657 | - |
| Tetherin | *Rousettus aegyptiacus* | Egyptian fruit bat | MT274401 | - | PRJNA300284 | - |
| Tetherin | *Tadarida brasiliensis* | Mexican free-tailed bat | MT274402 | - | PRJNA184055 | - |
| Tetherin | *Taphozous melanopogon* | Black-bearded tomb bat | MT274403 | - | PRJNA255191 | - |
| Tetherin | *Uroderma bilobatum* | Tent-making bat | MT274404 | - | PRJNA268573 | - |
| **Other mammals** |  |  |  |  |  |  |
| Tetherin A Isoform A | *Bos taurus* | Cow | - | J7M5G6 | - | - |
| Tetherin A Isoform B | *Bos taurus* | Cow | - | J7MAQ7 | - | - |
| Tetherin B | *Bos taurus* | Cow | - | J7M2B2 | - | - |
| Tetherin | *Felis catus* | Cat | - | F8R0X8 | - | - |
| L-tetherin | *Homo sapiens* | Human | - | Q10589-1 | - | - |
| S-tetherin | *Homo sapiens* | Human | - | Q10589-2 | - | - |
| Tetherin | *Loxodonta africana* | Elephant | - | G3UKK9 | - | - |
| Tetherin | *Macaca mulatta* | Rhesus Macaque | - | C4P4A2 | - | - |
| Tetherin | *Mus musculus* | Mouse | - | Q8R2Q8 | - | - |
| Tetherin A | *Ovis aries* | Sheep | - | D5JZS7 | - | - |
| Tetherin B | *Ovis aries* | Sheep | - | D5JZS8 | - | - |
| Tetherin | *Pan troglodytes* | Chimpanzee | - | D7RVC2 | - | - |

The dual-tyrosine Y·x·Y motif, which is critical for mediating viral-particle endocytosis and is involved in immune signalling (31, 40), is variable among bat species and exists in different combinations of Y|C·x·Y|H (Figure 1A and 1C). All bat species possess at least one tyrosine residue within this motif (Figure 1C). Conservation of the protein domain organization and key structural motifs of bat tetherin, despite significant amino acid sequence diversity, supports the present understanding of tetherin as a protein whose functions are mediated through structural rather than sequence-mediated interactions (28).

### Bat tetherin genes are under significant positive selection

To assess the evolutionary pressure applied during the speciation of bats, a selection test was performed to analyse bat tetherin genes spanning to ∼65 million years ago (mya) of Chiropteran history. First, to avoid the inclusion of paralogs in the selection test, a phylogenetic analysis of the tetherin sequences was performed (Figure 2). Based on branch lengths and branching patterns, *Myotis macropus* tetherin 2A (BST-2A) was determined to be the ortholog of tetherin in other bat species. *M. macropus* tetherins B-E were considered to represent paralogs and excluded from the subsequent selection test. The selection test revealed that overall, bat tetherin genes have been subjected to positive selection, with an average ratio of non-synonymous (dN) to synonymous (dS) mutations, dN/dS, of 1.125 over the Chiropteran phylogeny, with numerous specific positions subjected to a large degree of positive selection, with dN/dS values of 2.497 - 2.561 (Table 2). These sites are predominantly located in the transmembrane domain and the regions of the cytoplasmic and extracellular domains immediately adjacent to the transmembrane domain (Figure 1A). This analysis suggests the presence of evolutionary pressure exerted on tetherin, by viral antagonists or countermeasures that target residues within tetherin, following the speciation of the Chiropteran order. The apparent ubiquity of tetherin among bats, in concert with significant sequence diversity and evidence of strong positive selection pressures operating on each species, supports the notion that tetherin plays a major role in the antiviral repertoire of bats.

**Figure 2.**
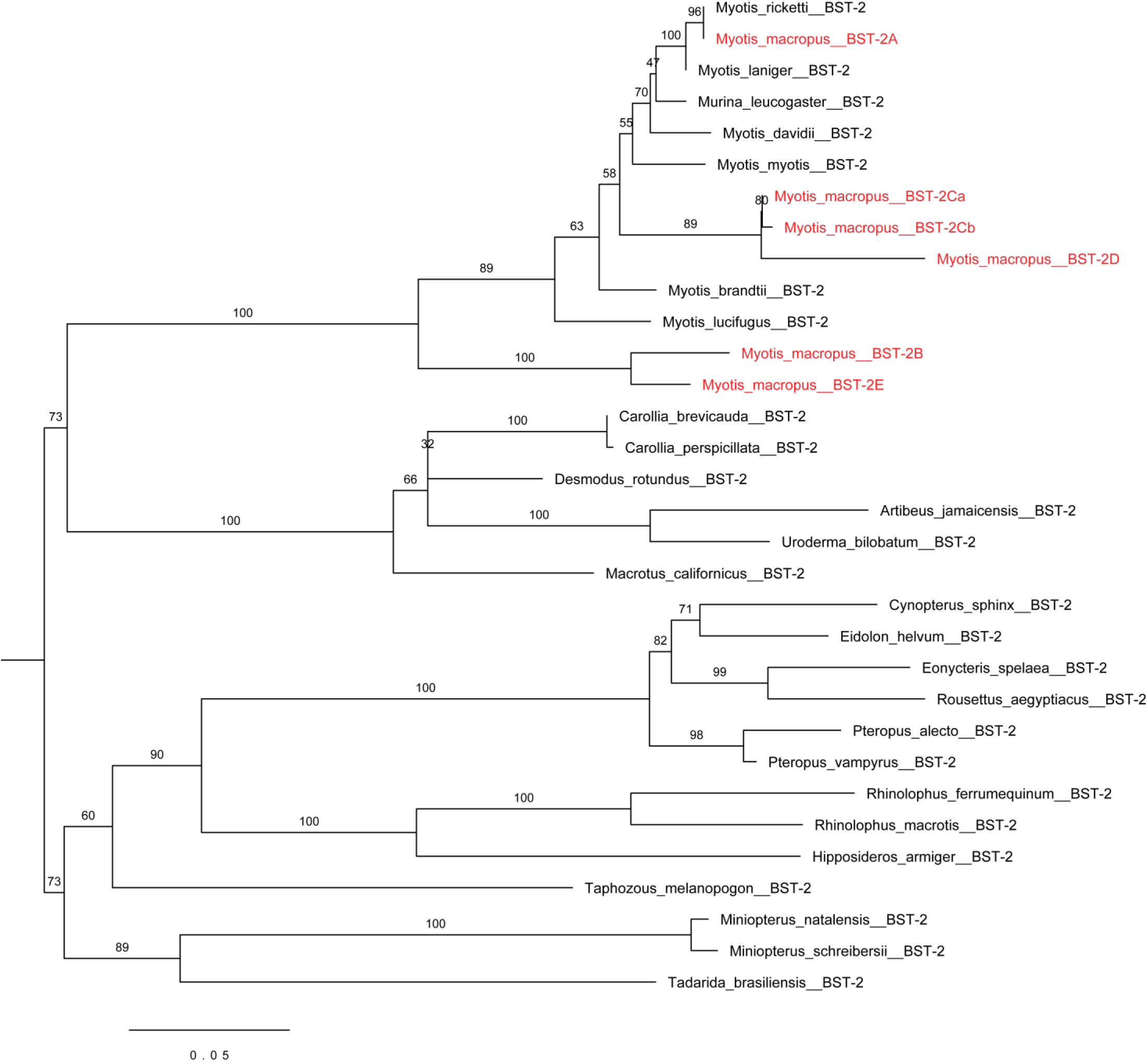
Maximum likelihood phylogenetic analysis of bat tetherin (BST-2) sequences. The phylogeny was generated with CODEML implemented in PAML4 software (74). The input tree was generated based on phylogeny generated by TimeTree (75). Nodes are labelled with the name of the bat species from which the tetherin sequences are derived. Multiple tetherin paralogs are represented for *Myotis macropus*, labelled in red. The scale represents the number of nucleotide substitutions per site.

**Table 2.**
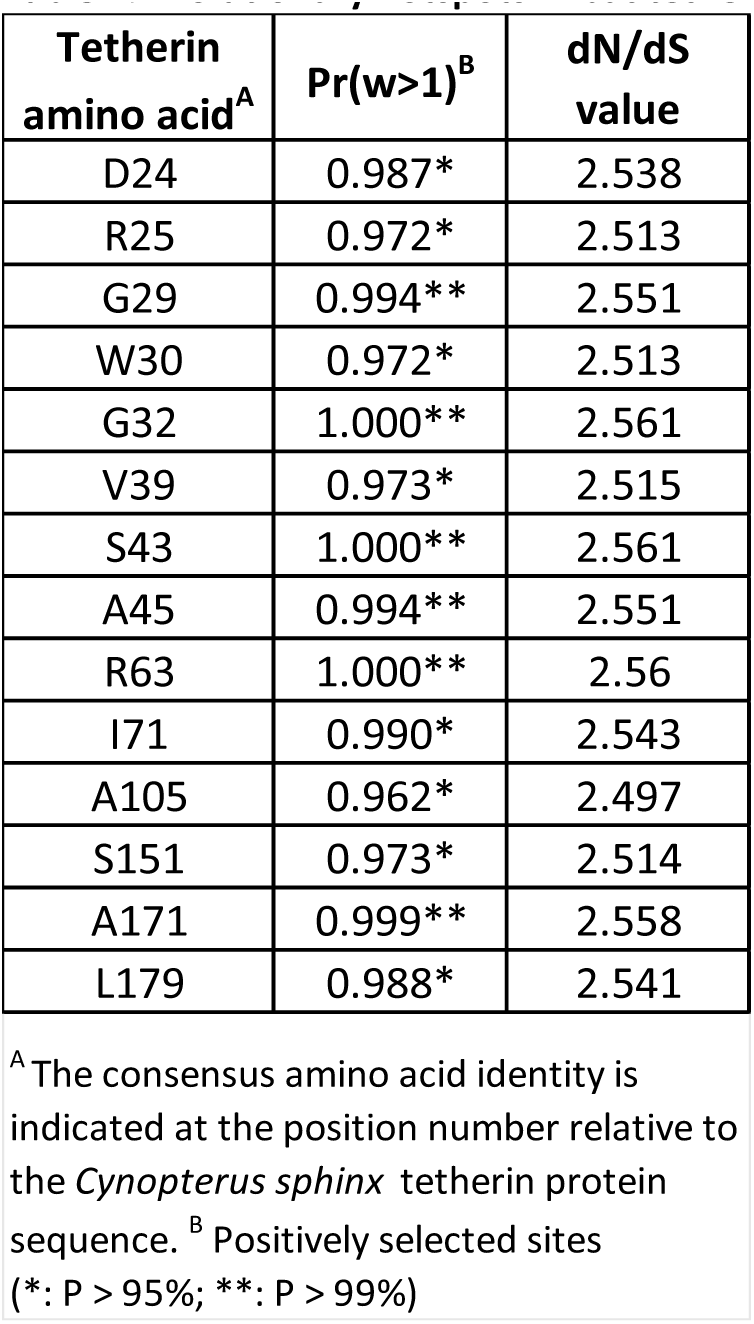
Evolutionary hotspots in bat tetherin

### Fruit bats possess unique structural isoforms of the tetherin protein generated through alternative splicing of a single tetherin gene

The expression of the fruit bat tetherin gene was initially assessed by a BLASTn search within the transcriptome of an Australian fruit bat, the Black flying fox, *Pteropus alecto*, using the human tetherin amino acid sequence as the search query. We identified 30 contigs matching the query (lowest E-value = 9.21 x 10^-23^). A single *P. alecto* homolog of tetherin, herein described as isoform A, homologous to human l-tetherin and the predicted X1 isoform (GenBank: XM_006904279), was identified. *P. alecto* tetherin isoform A has low primary amino acid sequence conservation (37%) compared to human l-tetherin (Figure 3A) and other mammalian (28 - 47%) tetherin sequences homologous to human l-tetherin (Figure 3B). Despite the low amino acid sequence identity, the predicted secondary structures and protein domains are largely conserved between bats and other mammals (Figure 3B). All previously reported tetherin proteins containing the cytoplasmic (CD), transmembrane (TM), extracellular domains (ED), and the post-translationally cleaved GPI-anchor signal peptide (GSP) with the exception of sheep tetherin B, which does not contain the GSP domain.

**Figure 3.**
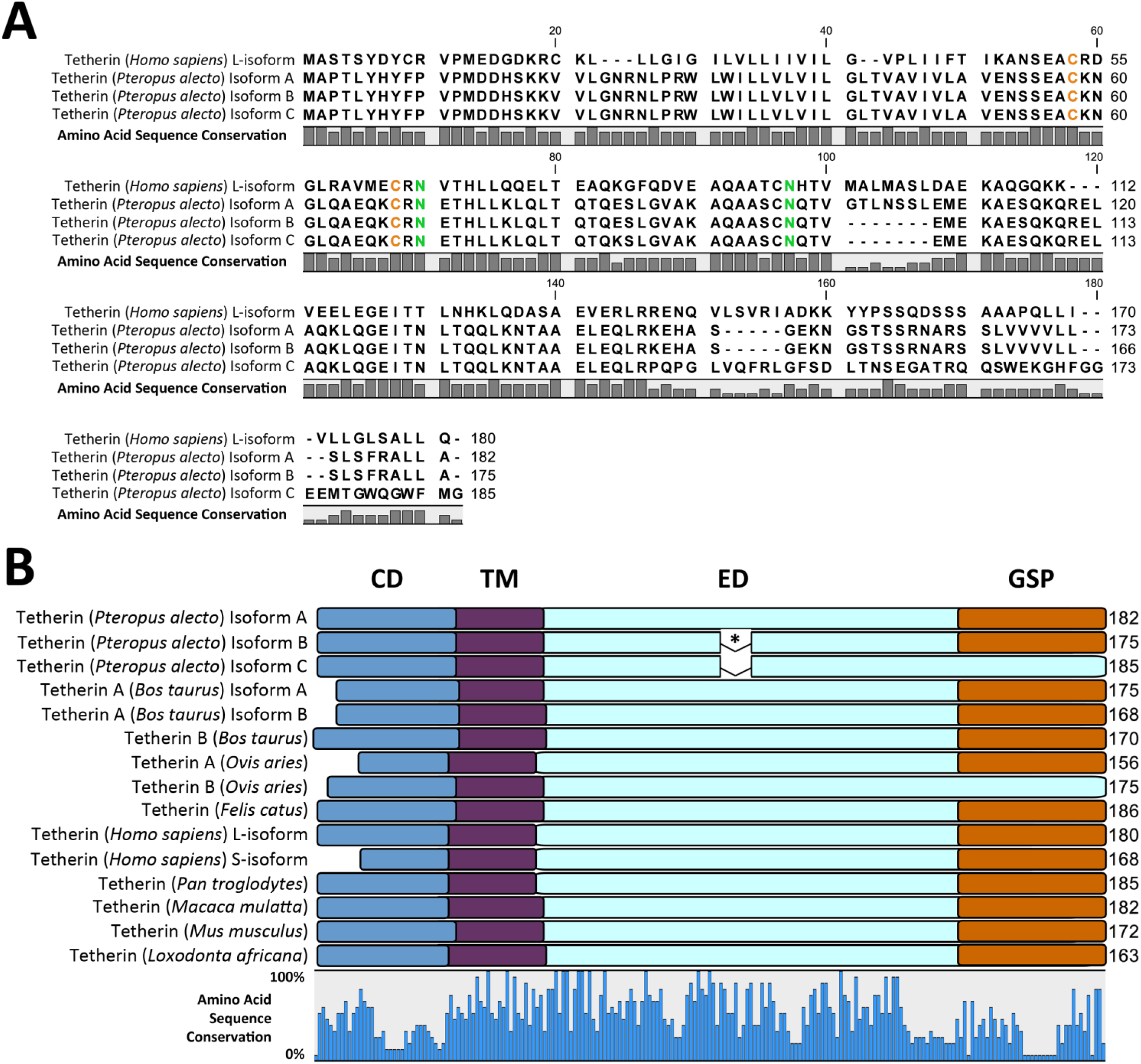
Multiple sequence alignment (MSA) and protein domains of mammalian tetherin amino acid sequences. **A.** MSA reveals amino acid residues differing between human and *P. alecto* sequences which are indicated by the amino acid sequence conservation bar graph below the sequences. Conserved cysteine and asparagine residues are coloured orange and green, respectively. **B.** The protein domains of mammalian tetherin proteins are depicted as overlays of an MSA. Sequence conservation is indicated by the bar graph below the alignment. Protein domains are colour coded: CD (blue), cytoplasmic domain; TM (purple), transmembrane domain; ED (light blue), extracellular domain; GSP (orange), glycophosphatidylinositol signal peptide; * a 7 AA exclusion in *P. alecto* tetherin isoforms B and C relative to isoform A is indicated.

To verify the expression of bat tetherin isoform A, a cDNA library was prepared from *P. alecto* spleen tissue, comprising several immune cell types, and assessed for the presence of transcripts matching that identified in the *P. alecto* transcriptome. Primers were designed to flank the open reading frame of tetherin isoform A, and amplicons were generated by PCR. Amplicons were then cloned into plasmid vectors for DNA sequencing. This analysis revealed two additional isoforms of tetherin, isoforms B & C. Isoform B was homologous to the computationally predicted isoform X2 (GenBank: XM_015587122). Both isoforms B and C were predicted to contain structural differences relative to isoform A. They both possess a 7 AA exclusion in the middle of the extracellular domain while isoform C was additionally predicted to contain an alternative C-terminus, without the GSP domain (Figure 3B). The absence of a GSP domain suggests that it is unlikely that parallel homodimers of isoform C would possess the capacity to tether viral particles to the cell.

Mapping the tetherin cDNA sequences against the *P. alecto* genome identified a single tetherin gene located on Scaffold KB030270 (Figure 4A), between the MVB12A (multivesicular body subunit 12A) and PLVAP (plasmalemma vesicle associated protein) genes, similar to the human tetherin gene (Viewable in the Ensembl genome browser: http://www.ensembl.org/Homo_sapiens/Location/View?db=core;g=ENSG00000130303;r=19:17402886-17405683). *P. alecto* Isoform B and C mRNA are generated from the use of an alternative splice acceptor site for the first intron, resulting in the 7 AA exclusion, while the distinct C-terminus of isoform C is a result of alternative mRNA splicing that incorporates exon 5 while excluding exon 4 used in isoform A and B (Figure 4B).

**Figure 4.**
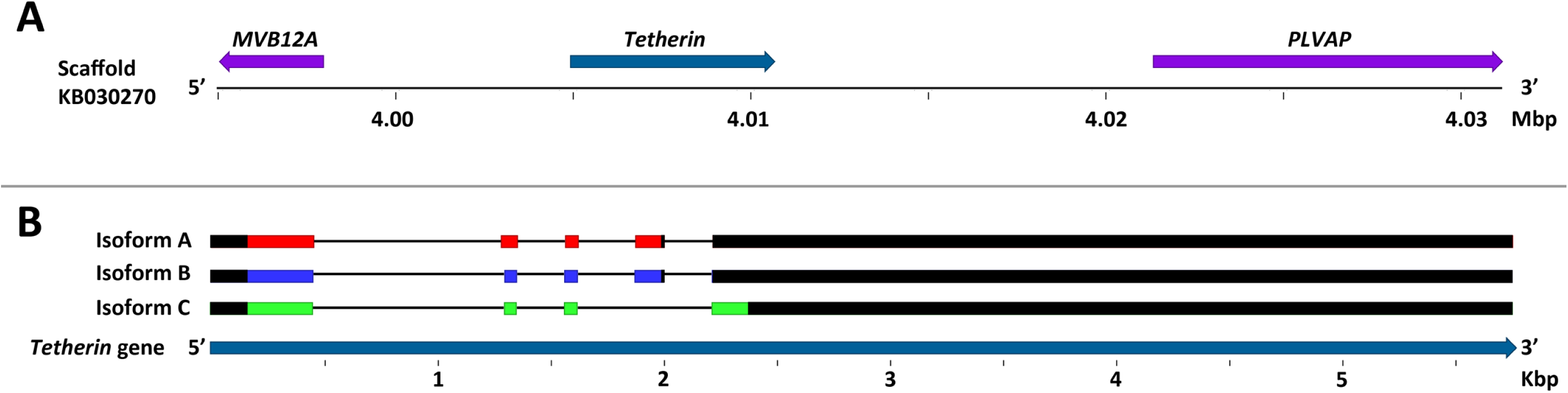
Mapping the tetherin gene to the *P. alecto* genome. **A.** The tetherin gene location (dark blue) and neighbouring genes (purple). Arrows indicate the orientation of each gene’s protein coding domain sequence. Sequence numbers and gene orientations are relative to the beginning of the gene scaffold in the 5’ to 3’ direction. **B.** The tetherin mRNA exons are depicted and colour coded: Isoform A, red; Isoform B, blue; Isoform C, green. Exons are mapped to scale against the tetherin gene. Sequence numbers are relative to the beginning of the gene. The 5’ and 3’ untranslated regions of the mRNA are coloured black.

### Vesper bats possess multiple tetherin genes that contain unique structural variations

To further investigate the diversity of tetherin in non-fruit bat species, we amplified the tetherin nucleotide sequences for the vesper bat species *Myotis ricketti* and *Miniopterus schreibersii* using primers designed on the basis of the transcriptome sequences (Table 3A). These primers were used for PCR of cDNA samples generated from *M. ricketti* spleen tissue and a *M. schreibersii* kidney cell line to amplify the tetherin coding domain sequences (Table 3A). Primers for *M. ricketti* were also used to amplify the tetherin coding region from cDNA generated from a *Myotis macropus* kidney cell line as it was reasoned that since these species belong to the same genus, primers designed for one might be capable of amplifying tetherin from the other. Following PCR, amplified DNA from *M. ricketti, M. macropus*, and *M. schreibersii*, were gel purified, cloned and sequenced.

**Table 3.**
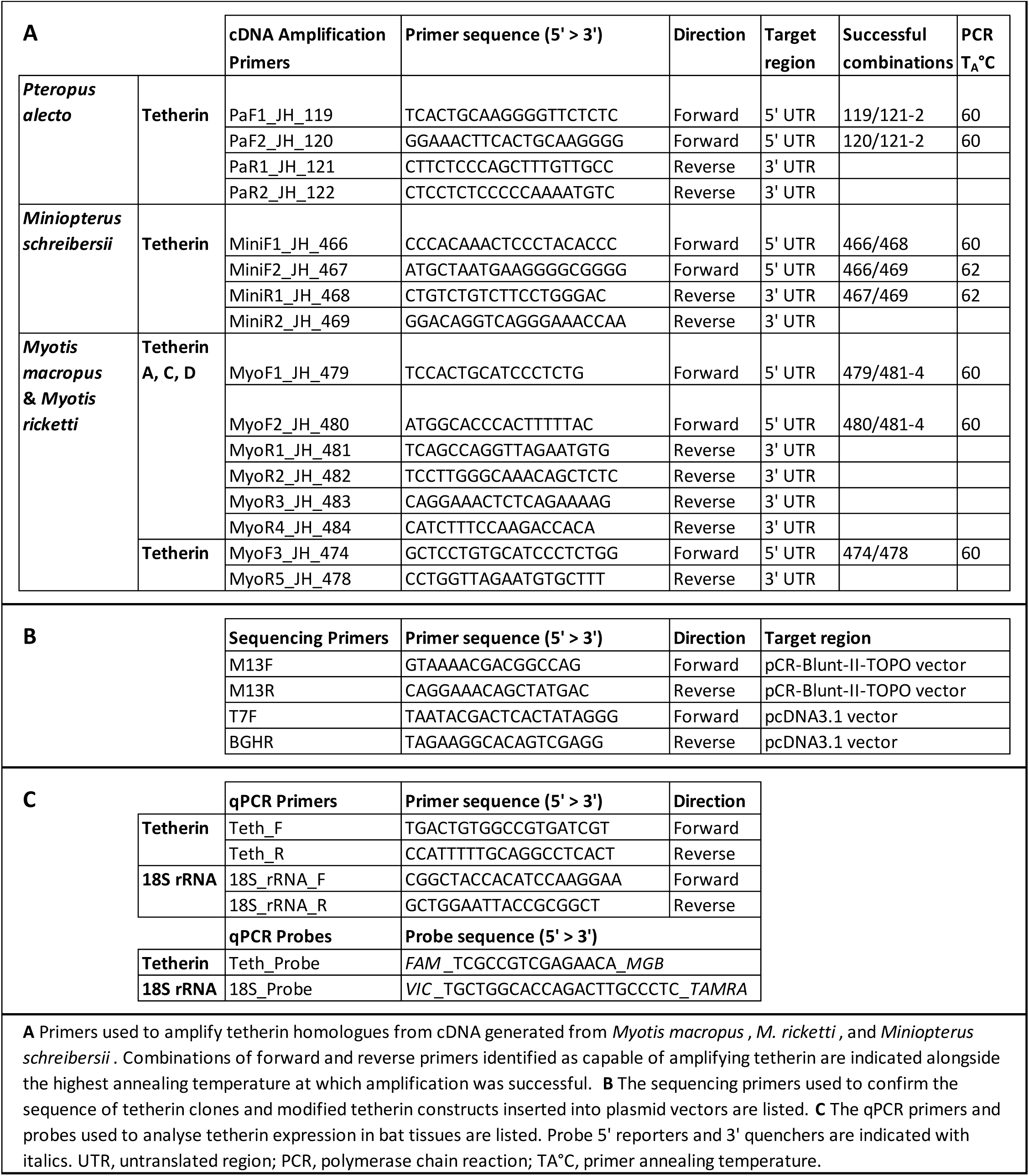
cDNA amplification and vector sequencing primers for fruit bat and vesper bat tetherins.

These microbats were found to express various homologs of tetherin that include encoding of unique structural variations (Figure 5A and 5B). The tetherin homolog of human l-tetherin, predicted for *M. schreibersii,* was detected (Figure 5A). Five unique tetherin variants, differing in their encoded amino acid sequences (tetherin A – E), were identified for *M. macropus*, two of which, tetherin A and B, were also detected in *M. ricketti* (Figure 5A). These tetherin variants encode three distinct homologs (tetherin A, C, D) of the human l-tetherin, sharing the same protein domains but differing in their amino acid sequence (Figure 5).

**Figure 5.**
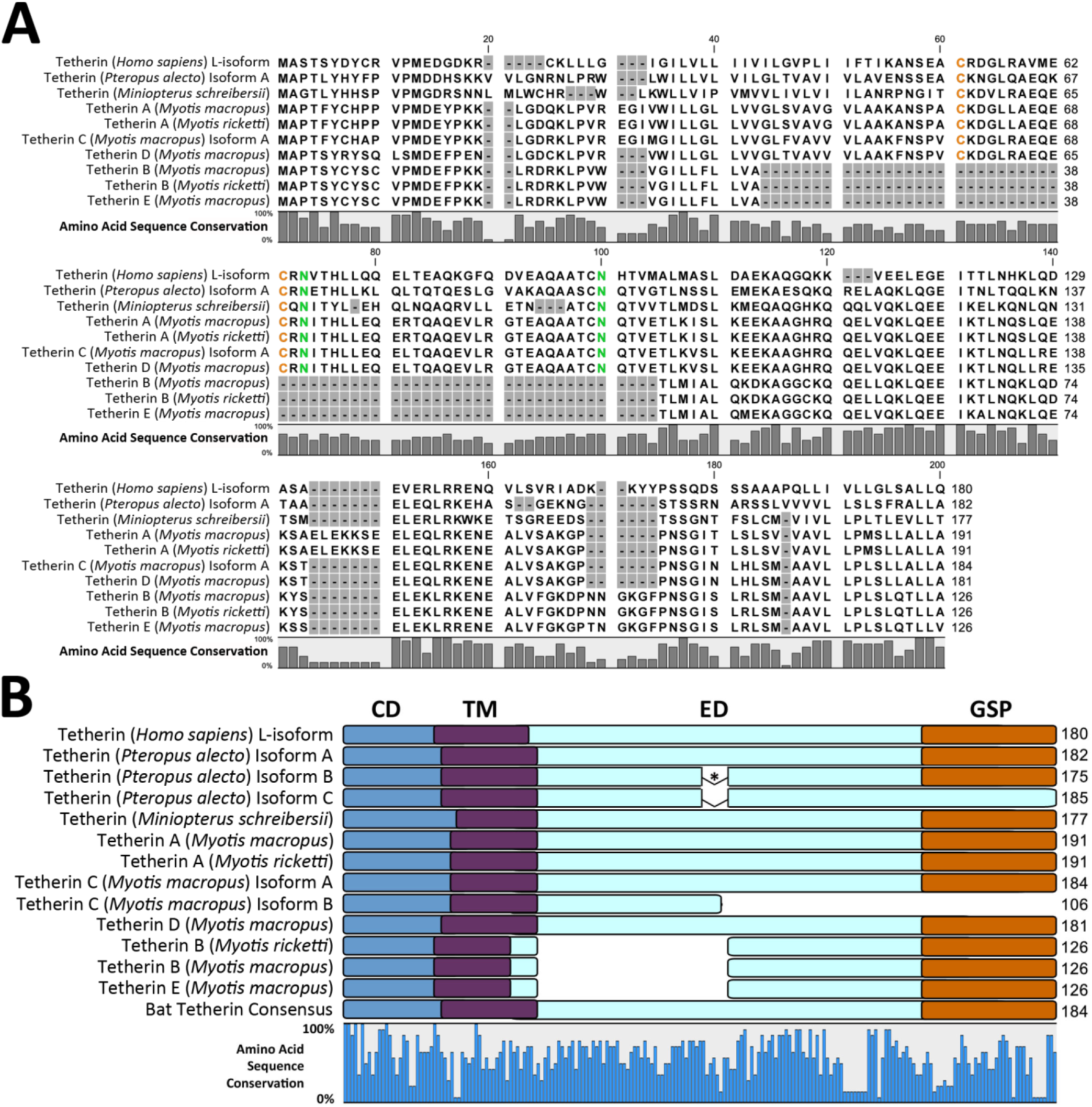
Multiple sequence alignment (MSA) and protein domains of fruit bat and vesper bat tetherin amino acid sequences. **A.** The percentage of sequence conservation is indicated by a bar plot below the MSA. Gap positions are shaded in grey. Conserved cysteine and asparagine residues are coloured orange and green, respectively. **B.** The protein domains of bat tetherin proteins are depicted as overlays of an MSA. Amino acid sequence conservation is indicated by the bar graph below the alignment. Protein domains are indicated and colour coded as follows: CD (blue), cytoplasmic domain; TM (purple), transmembrane domain; ED, (light blue) extracellular domain; GSP (orange), glycophosphatidylinositol signal peptide. * The 7 AA exclusion in *P. alecto* tetherin isoform B and C relative to isoform A is indicated.

An additional splice variant (isoform B) of *M. macropus* tetherin C was also identified. *M. macropus* tetherin C isoform B is predicted to have a C-terminal truncation and possesses only the cytoplasmic domain, transmembrane domain, and less than half of the extracellular domain (Figure 5B). Accordingly, it is predicted to lack the GPI signal peptide required for the post-translational addition of a GPI-anchor, also found for *P. alecto* tetherin isoform C (Figure 3B). The absence of a GPI anchor indicates that it is unlikely that homodimers of *M. macropus* tetherin C isoform B would possess the capacity to tether viral particles to the cell. *M. macropus* tetherin B and E were found to be structurally unique, possessing a large deletion of ∼60 amino acids in the extracellular coiled-coil domain (Figure 5B), compared to the other tetherin homologs, which has been shown to be critical for viral particle restriction (28, 43). This deletion results in the exclusion of the conserved disulphide bond-forming cysteine residues, and the conserved asparagine-linked glycosylation sites, indicating that this form of tetherin is unlikely to form dimers or be glycosylated in the manner of human tetherin (28). They are however predicted to possess all other tetherin domains including the transmembrane domain and GPI signal peptide, indicating that they may still be capable of tethering viral particles. *M. macropus* tetherin B and E share 92.1% amino acid sequence identity indicating they are the product of a relatively recent gene duplication.

While there is no available genome for *M. macropus*, the most closely related publicly accessible genome assemblies are for the relatively distantly related *Myotis lucifugus* and *M. davidii,* which diverged from *M. macropus* approximately 19 and 14 mya, respectively (44, 45). For comparison, humans and chimpanzees diverged approximately 6.5 mya (46). A BLASTn analysis of the *M. macropus* tetherin nucleotide sequences against the *M. davidii* and *M. lucifugus* genomes revealed first that both species were too distantly related to *M. macropus* to match each *M. macropus* gene product specific to *M. davidii* or *M. lucifugus* gene regions. However, the *M. lucifugus* genome assembly included a single gene scaffold (GL430608) which contained matches to all query *M. macropus* tetherin sequences, comprising seven potential *M. lucifugus* tetherin genes (Figure 6A) and indicating the presence of a tetherin gene locus within this scaffold. Further analysis of the mapped exons, as described below, enabled confirmation of the *M. lucifugus* tetherin regions. The *M. lucifugus* tetherin gene locus was located between the MVB12A and PLVAP genes, as observed for the single *P. alecto* tetherin gene (Figures 4 and 6).

**Figure 6.**
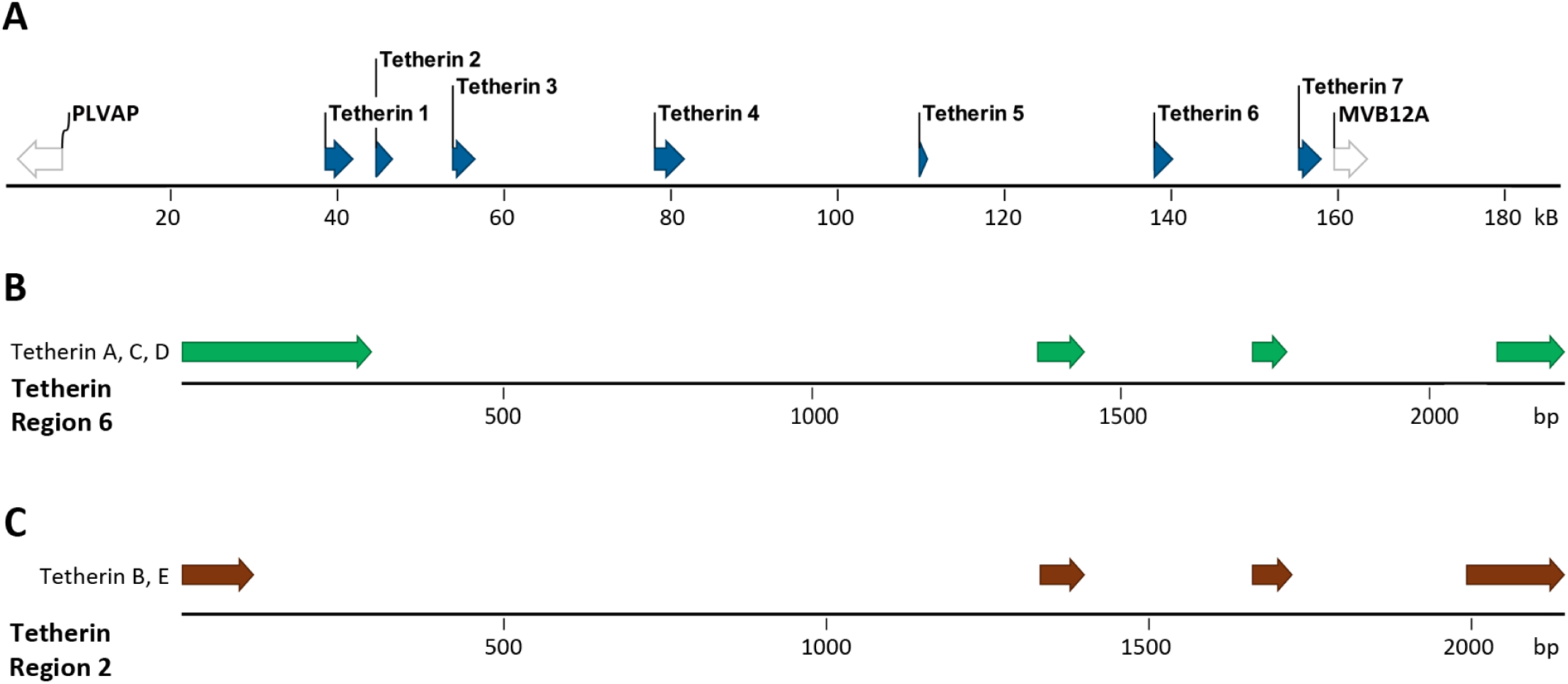
Mapping the tetherin genes of *M. macropus* to the *M. lucifugus* genome. Tetherin sequences derived from *M. macropus* cDNA mapped against the publicly available genome of *M. lucifugus* (Myoluc2.0 genome assembly). **A.** The tetherin gene locus of *M. lucifugus* is located within scaffold GL430608 flanked by the PLVAP and MVB12A genes. **B.** *M. macropus* tetherin A, C, and D mapped against *M. lucifugus* tetherin region 6. **C.** *M. macropus* tetherin B and E mapped against *M. lucifugus* tetherin region 2.

The genome assembly of *M. davidii* also contained significant matches to *M. macropus* tetherin sequences; however, these were spread across seven different gene scaffolds against which no *M. macropus* tetherin coding domain sequence could be matched in its entirety. For this reason, further genomic analysis of *Myotis* tetherin genes was confined to the *M. lucifugus* genome.

The *M. lucifugus* tetherin gene locus (Figure 6A) contains seven potential genes that are defined by the presence of two or more exons mapping to the *M. macropus* tetherin cDNA. Three exons across tetherin gene regions 1, 2, and 6, possess all of the coding exons required to express homologs to *M. macropus* tetherin A, C, and D (Figure 6B). Furthermore, *M. lucifugus* tetherin gene regions 2, and 3 possess all of the coding exons required to produce homologs to *M. macropus* tetherin B and E (Figure 6C). These data indicate that microbats of the genus *Myotis* possess a greater number and diversity of tetherin genes than any mammal reported to date.

These findings expand upon a recent report by Hölzer and colleagues (18), that identified the expression of three tetherin paralogs, upregulated by interferon treatment, in the vesper bat, *M. daubentonii*. Mapping of these paralogs against the genome of *M. lucifugus* revealed four tetherin genes, currently listed as novel gene predictions within the Ensembl database: ENSMLUG00000023691, ENSMLUG00000029243, ENSMLUG00000026989, and ENSMLUG00000023562 (18). Herein these genes are labelled Tetherin 1, 2, 6, and 7, respectively (Figure 6A).

### Tetherin expression pattern in *P. alecto* tissue samples

Tetherin is widely and variably expressed across human tissues and is upregulated in response to type I interferon (35, 36). To assess the relative expression of bat tetherin across different tissue types, tissue samples were obtained from three individual *P. alecto* bats: a male, a pregnant female, and a juvenile male. Using primers that amplify all three isoforms of *P. alecto* tetherin, expression was analysed by qPCR with Ct values normalised against the 18S rRNA housekeeping gene (Figure 7). The two male bats exhibited similar tetherin expression patterns, with greatest levels in the thymus. A thymus tissue sample was not obtained from the female, although a similar pattern to the male bats was observed among most other tissues. The female bat had higher expression in the lung compared to the male bats, possibly reflecting an active response to a respiratory infection, an artefact of hormonal regulation, or natural heterogeneity between bats.

**Figure 7.**
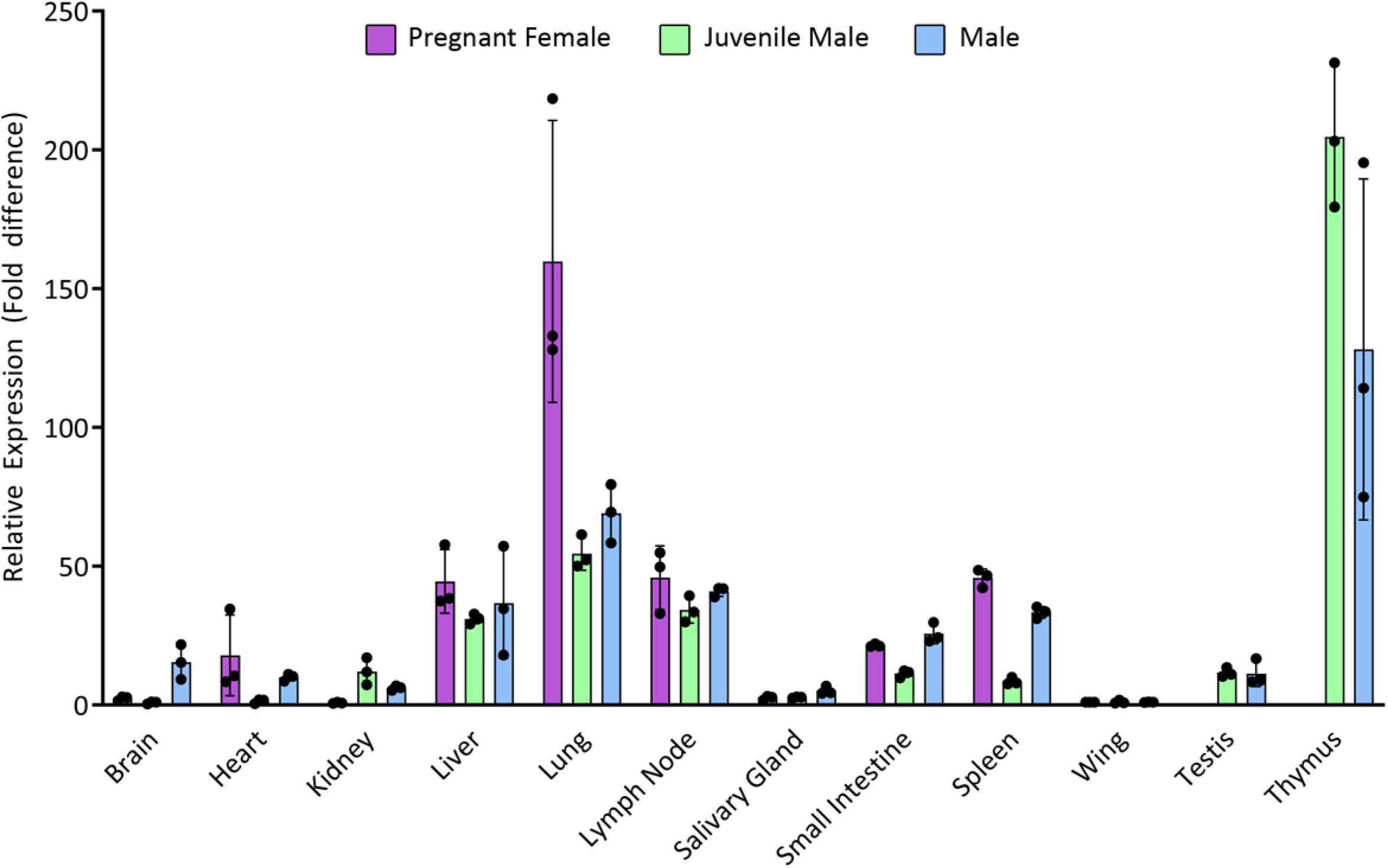
Relative tetherin expression across tissues from *P. alecto*. Expression of tetherin mRNA assessed by qRT-PCR analysis of tissues from three bats. In all cases, cycle threshold (Ct) values for tetherin were normalised against expression of the 18S rRNA gene. For all individuals, relative tetherin expression in each tissue is calibrated against expression in wing tissue. A thymus sample was not obtained for the female bat. Error bars represent the standard error (N = 3).

Previous studies in vesper bats have demonstrated that tetherin is upregulated in response to type I interferon alpha stimulation (18). In *P. alecto*, lipopolysaccharide (LPS) treatment increases the percentage of B-cells in the spleen (47), and polyinosinic:polycytidylic acid (PIC) treatment is observed to increase the expression of the interferon-stimulated gene, *ISG54* (48). To determine if pathogen-mediated stimulation can upregulate tetherin expression in fruit bats we analysed the transcriptome of *P. alecto* spleen tissue from animals treated *in vivo* with the Toll-like receptor (TLR) agonists, LPS or PIC. We found that *P. alecto* tetherin was significantly upregulated by LPS (P = 0.028) and PIC (P = 0.004) (Figure 8A).

**Figure 8.**
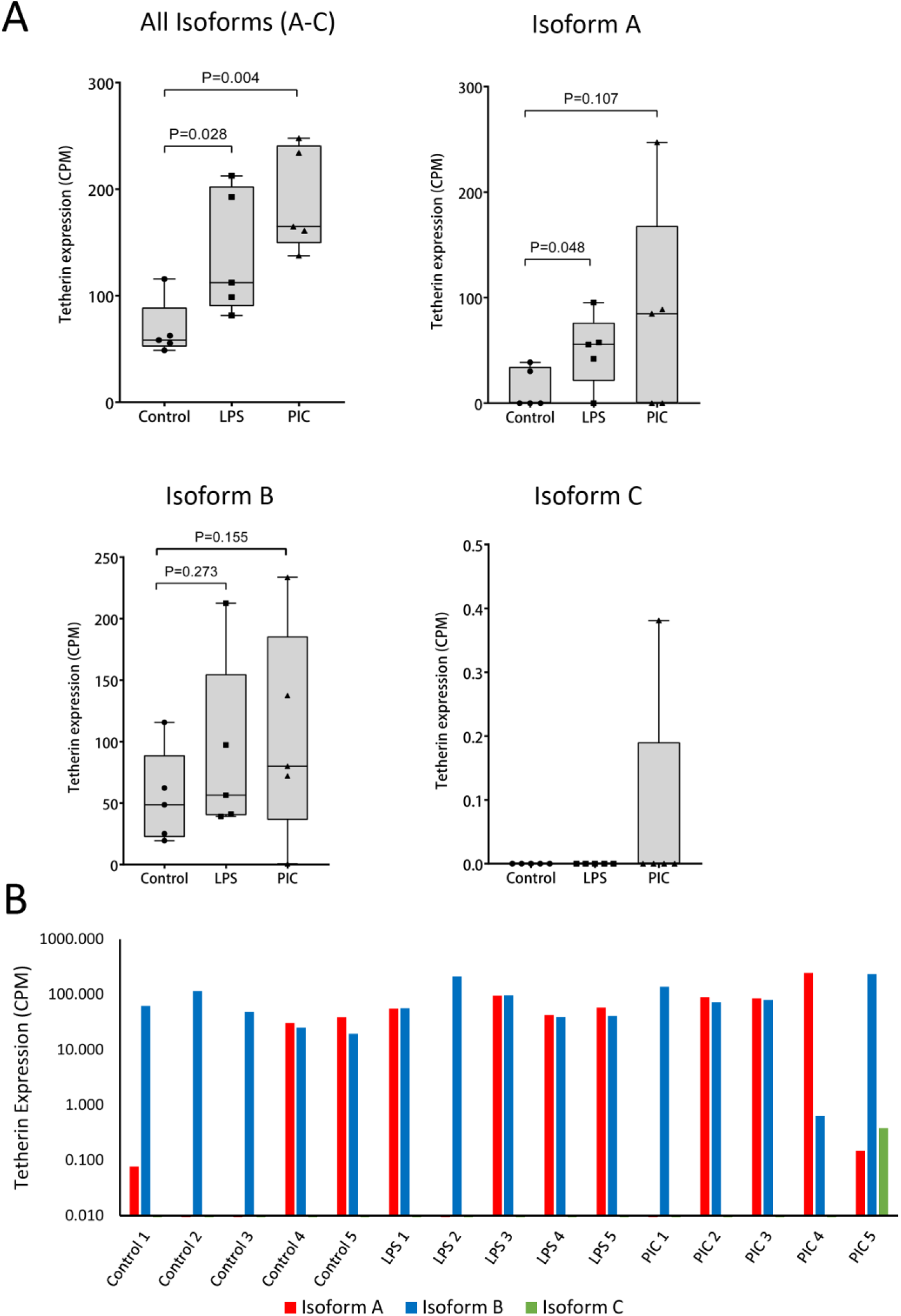
Expression of alternative isoforms of tetherin in stimulated and unstimulated *P. alecto* spleen tissue. Spleen tissue from 15 individuals were unstimulated, or stimulated with lipopolysaccharide (LPS) or polyinosinic:polycytidylic acid (PIC) (N = 5 for each treatment). Tetherin expression was determined from RNA-Seq sequence reads mapped against the *P. alecto* genomic scaffold, KB030270, which harbours the tetherin gene. Tetherin expression levels are represented as normalised counts-per-million (CPM) reads. **A.** Tetherin expression by isoform. P-values were determined using the one-tailed Mann-Whitney test. **B.** Expression of tetherin isoform in each individual sample.

We next assessed if there was a difference in the expression of the alternative tetherin isoforms A, B and C and the impact on the expression of each isoform following treatment with the TLR agonists LPS and PIC. This analysis revealed that tetherin is predominantly expressed as isoform B (14/15 samples; Figure 8B) while stimulation with LPS significantly increased the expression of Isoform A (P = 0.048), but not Isoform B or C (Figure 8A). Stimulation with PIC increased the expression of isoform A and B compared to unstimulated tissue; however, this effect was variable among individual bat spleen samples and did not reach significance. Expression of Isoform C was only observed in a single PIC treated spleen sample. These data show that tetherin isoforms are expressed in tissue and can be upregulated by stimulation with TLR-agonists.

### *P. alecto* tetherin protein expression and cellular localisation

To characterise bat tetherin, we analysed tetherin from two Australian bats, the fruit bat *P. alecto* and the vesper bat *M. macropus*. The coding sequences of the *P. alecto* tetherin isoforms A, B, and C, and *M. macropus* tetherin A and B, were cloned into mammalian expression plasmids. While numerous vesper bat tetherins were identified in this study, *M. macropus* tetherin A and tetherin B were selected for further analysis as tetherin A is a homolog of human l-tetherin and tetherin B was the most structurally unique, possessing a large deletion within the extracellular coiled-coil domain compared to tetherin A (Figure 5B).

All tetherin coding sequences were modified with the addition of a haemagglutinin (HA) tag sequence (N-SG**YPYDVPDYA**GS-C) to enable antibody-based detection of protein expression. In all cases the HA tag was inserted at the position immediately following the end of the coiled-coil region of the extracellular domain (Figure 1A) which is the equivalent location to that previously utilised in the tagging of human tetherin (28).

To evaluate tetherin protein expression, mammalian HEK293T cells, which do not express tetherin in the absence of type I interferon stimulation (21), were transfected with bat tetherin expression constructs. Cell lysates were extracted using a method for enhanced extraction of GPI-anchored proteins (49) and tetherin expression was observed by non-reducing SDS-PAGE and Western blot analysis (Figure 9A and 9B). Predicted molecular weights of unglycosylated and GPI-anchored tetherin proteins are listed in Table 5. Human tetherin is glycosylated, and the *P. alecto* and *M. macropus* tetherins analysed here are predicted to contain several glycosylation sites by the GlycoMine prediction tool (50) (Table 6). For all isoforms of *P. alecto* tetherin and *M. macropus* tetherin A, the predicted glycosylation sites with the highest probability are the highly conserved N-linked glycosylation sites highlighted in Figures 3A and 5A. *M. macropus* tetherin B contains a 60 AA exclusion that spans both conserved glycosylation sites (Figure 5A); however it contains two additional predicted glycosylation sites with a probability score > 0.5, indicating that it is likely to be glycosylated at these sites. All predicted glycosylation sites with a prediction probability score > 0.5 are listed in Table 6.

**Figure 9.**
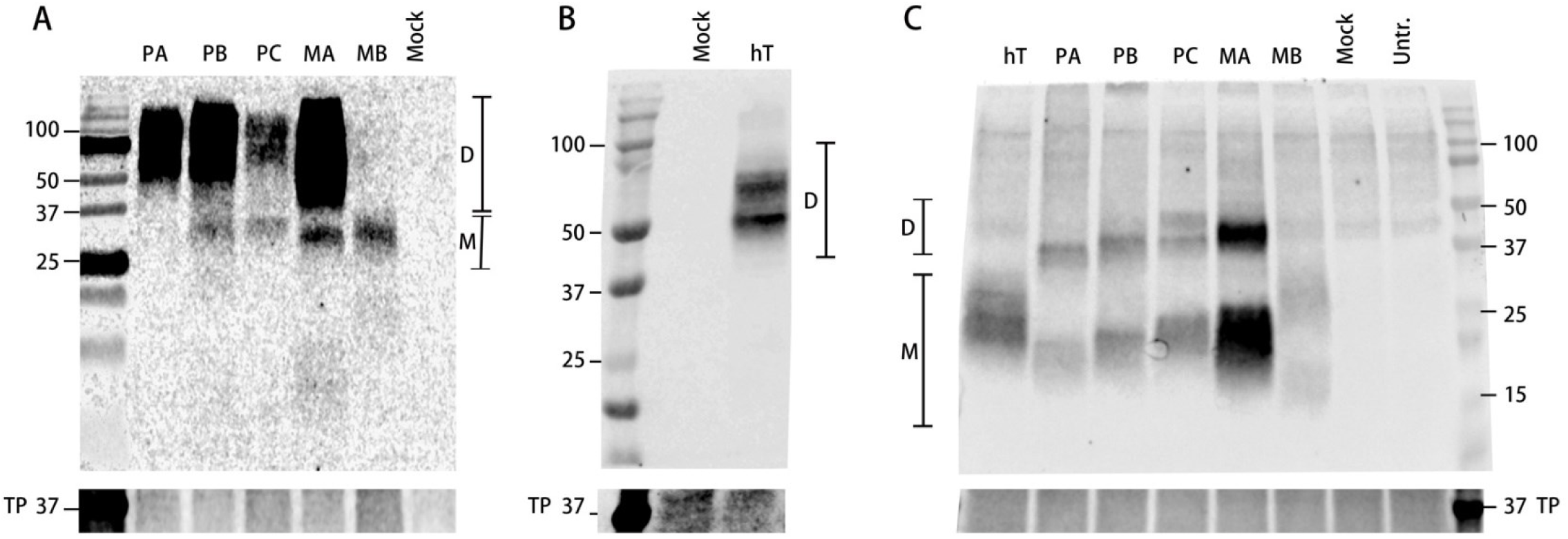
SDS-PAGE and Western blot analysis of the expression of modified bat tetherin constructs. Western blot analysis of the expression of haemagglutinin (HA) tagged *Pteropus alecto* tetherin isoforms A, B, C (PA, PB, and PC, respectively), and *Myotis Macropus* tetherin A and B (MA and MB, respectively), and human tetherin (hT) compared against and cells mock transfected (Mock) with an equivalent mass of empty vector plasmid pcDNA3.1, and untransfected (Untr.) cells. Tetherin was expressed in mammalian HEK293T cells and cell lysates were collected 48 h after transfection of cells with expression plasmids. **A.** Bat tetherin expression. SDS-PAGE of cell lysates run on a 12% polyacrylamide gel under non-reducing conditions. **B.** Human tetherin expression. SDS-PAGE of cell lysates run on an Any kD polyacrylamide gel (Bio-Rad) under non-reducing conditions. **C.** SDS-PAGE of cell lysates treated with PNGase to deglycosylate proteins and reduced in the presence of dithiothreitol and run on an Any kD polyacrylamide gel (Bio-Rad). Tetherin is present in dimeric (D) and monomeric (M) forms. Protein size scale is expressed in kDa. Total protein (TP) stain was used as a loading control.

**Table 4.**
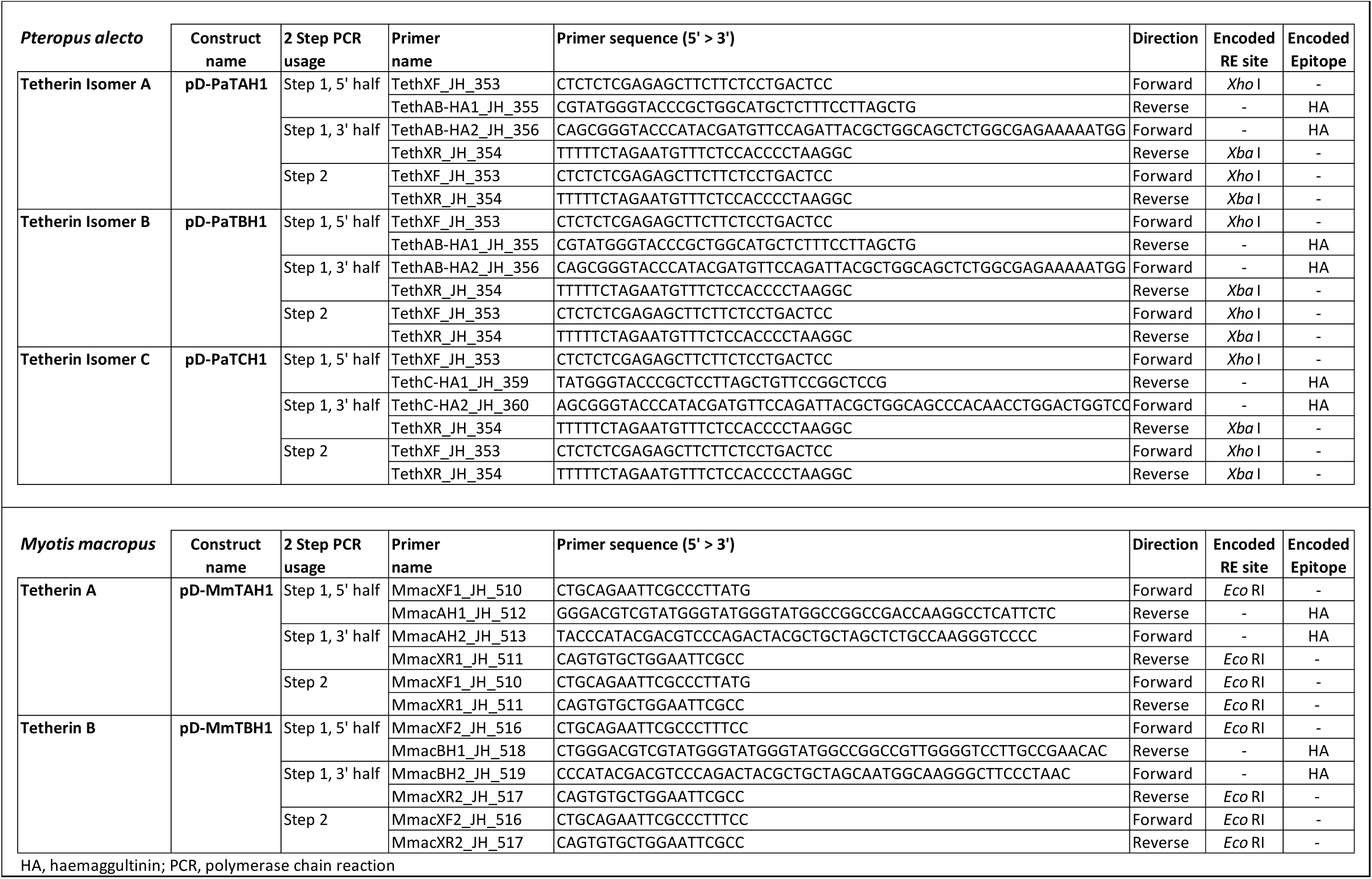
Primers used for the generation of haemagglutinin-tagged *Pteropus alecto* and *Myotis macropus* tetherin expression constructs through a 2-Step PCR process.

**Table 5.**
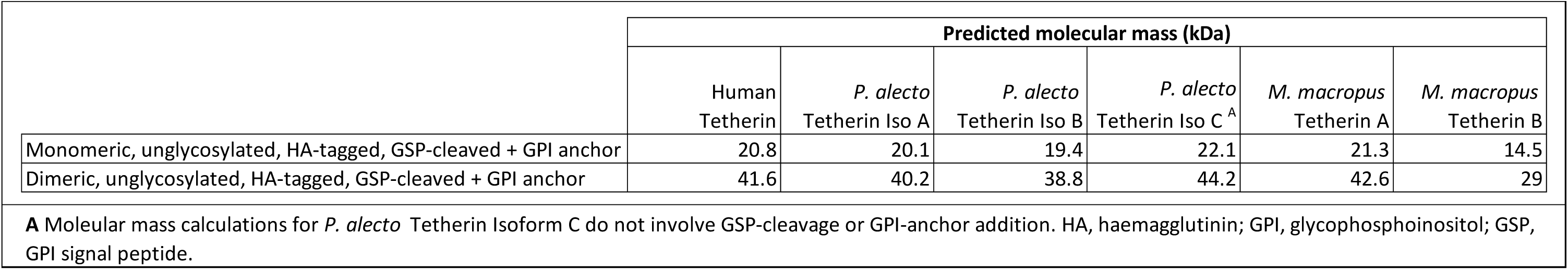
Predicted molecular mass for unglycosylated tetherin monomers and dimers.

|  | Predicted molecular mass (kDa) |  |  |  |  |  |
| --- | --- | --- | --- | --- | --- | --- |
|  | Human<br>Tetherin | *P. alecto*<br>Tetherin Iso A | *P. alecto*<br>Tetherin Iso B | *P. alecto*<br>Tetherin Iso C <sup>A</sup> | *M. macropus*<br>Tetherin A | *M. macropus*<br>Tetherin B |
| Monomeric, unglycosylated, HA-tagged, GSP-cleaved + GPI anchor | 20.8 | 20.1 | 19.4 | 22.1 | 21.3 | 14.5 |
| Dimeric, unglycosylated, HA-tagged, GSP-cleaved + GPI anchor | 41.6 | 40.2 | 38.8 | 44.2 | 42.6 | 29 |

**A** Molecular mass calculations for *P. alecto* Tetherin Isoform C do not involve GSP-cleavage or GPI-anchor addition. HA, haemagglutinin; GPI, glycosphosphoinositol; GSP, GPI signal peptide.

**Table 6.**
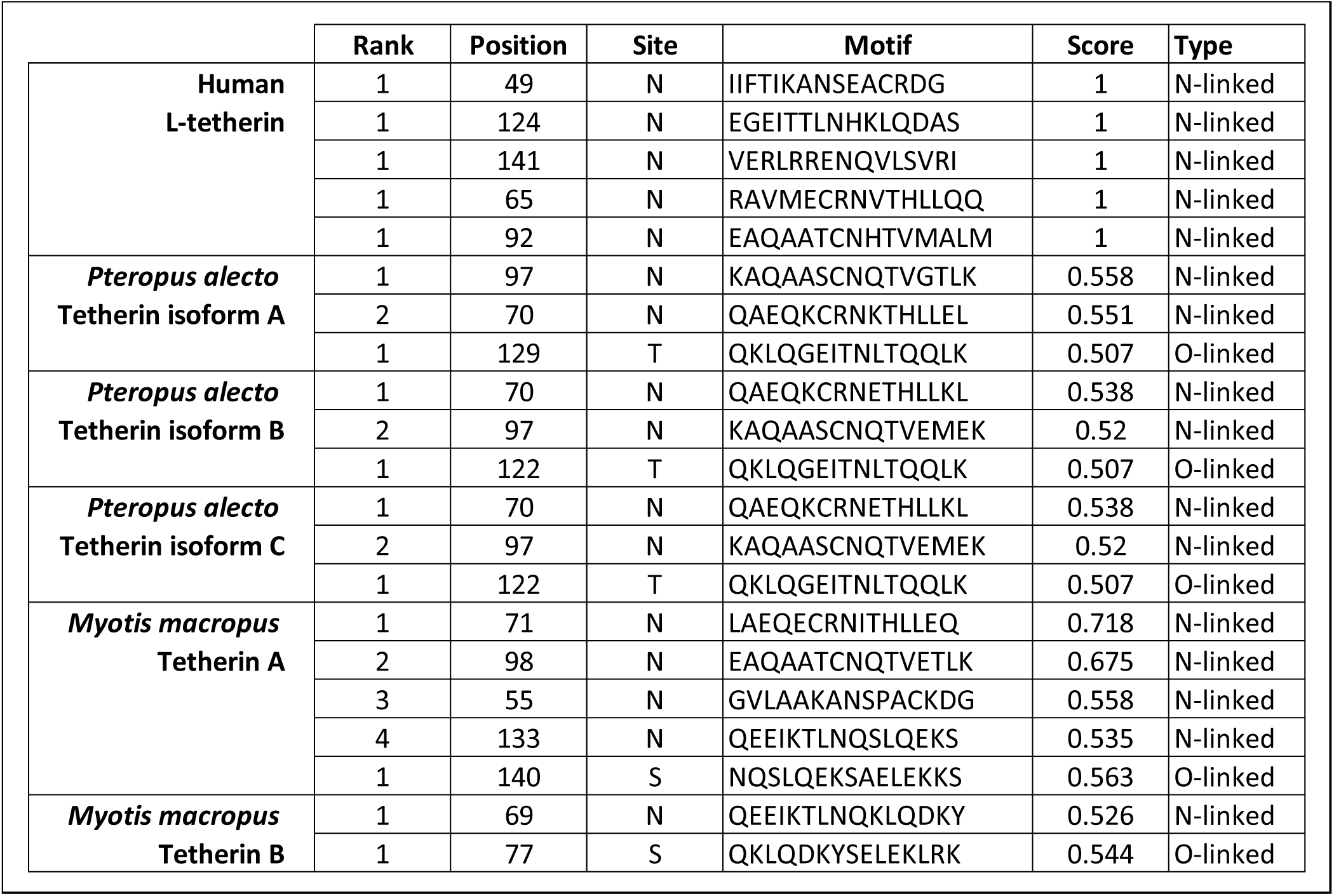
Predicted glycosylation sites with a probability score > 0.5, calculated by GlycoMine.

*P. alecto* tetherin isoforms A and B were detected primarily as broad bands at the expected positions for dimeric tetherin ∼56-75 kDa (Figure 9A), which is consistent with human tetherin (Figure 9B). *P. alecto* tetherin isoform C was present as both dimers and monomers (∼32-35 kDa), and some monomers were also detected for isoform B (Figure 9A). *M. macropus* tetherin A was also primarily detected as dimers, with some monomers also present (Figure 9A). In contrast, *M. macropus* tetherin B was detected only as a monomer (Figure 9A). The presence of broad/multiple bands in close proximity to each other likely reflects variable levels of tetherin glycosylation as previously reported for human tetherin (28). To better resolve bat tetherin we subjected samples to deglycosyation and a reducing agent (Figure 9C). Consistent with the expected effect, deglycosylated tetherins migrated at a lower molecular weight, ∼39-48 kDa for dimers and ∼15-24 kDa for monomers. Deglycosylation-treated, monomeric *M. macropus* tetherin B, migrated as two bands, the higher (∼30 kDa) corresponding to glycosylated monomers and the lower (∼15 kDa) matching the predicted molecular weight of unglycosylated tetherin (Table 5), indicating incomplete deglycosylation (Figure 9C). Interestingly, bat tetherin was resistant to reducing conditions, in contrast to human tetherin (Figure 9C), with only a partial decrease in bat tetherin dimers observed under conditions that almost fully reduced human tetherin to monomers. Because protein dimers may be stabilized by hydrophobic interactions, we analysed the hydrophobicity of the tetherin proteins *in silico*, using CLC’s hydrophobicity tool (Figure 10). These data indicated that both *P. alecto* tetherin and *M. macropu*s tetherin A were predicted to contain a more hydrophobic N-terminus (AA position 1-10) than human tetherin. This may account for the enhanced dimer stability observed here (Figure 9C). Together, these data show that HA-tagged tetherin can be expressed and that these proteins differ in their ability to dimerise.

**Figure 10.**
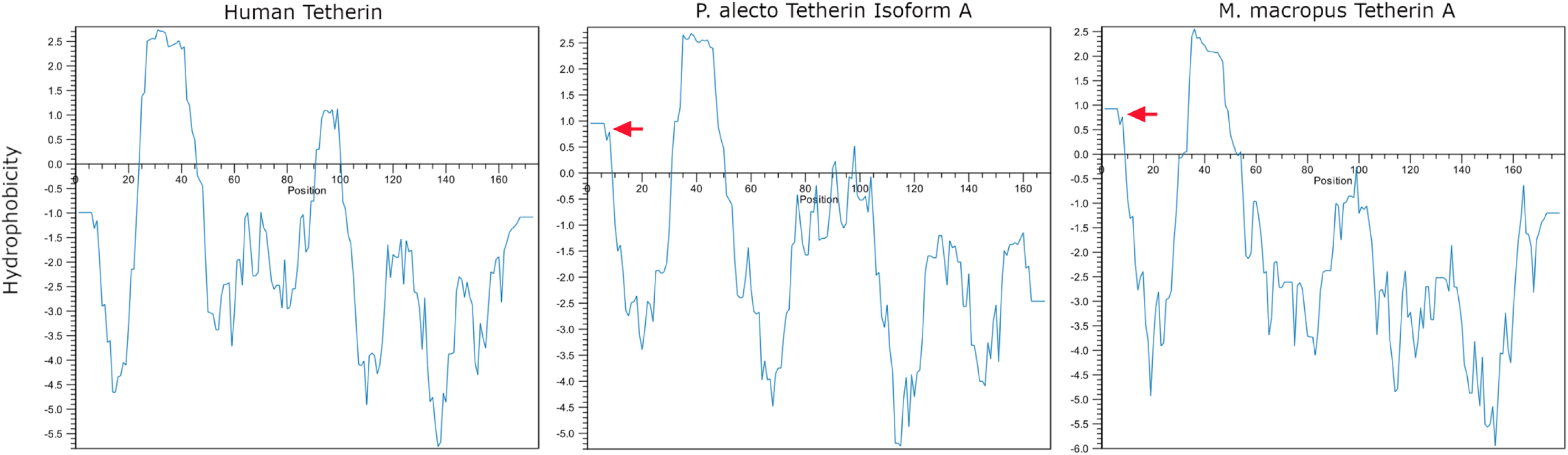
Hydrophobicity plot of human and bat tetherins. Amino acid sequences representing haemagglutinin-tagged, GSP-cleaved human tetherin, *Pteropus alecto* tetherin isoform A, and *Myotis macropus* tetherin A were analysed using the CLC hydrophobicity tool. Hydrophobicity was analysed across the sequence with an 11 AA sliding window. Positive hydrophobicity scores indicate hydrophobic regions. Red arrows indicate the small region of hydrophobicity at the N-terminus of the bat tetherins.

Human tetherin localises to the plasma membrane at sites of virus budding, in addition to membranes of the trans-Golgi network and recycling compartments (21, 51). To determine localisation of *P. alecto* tetherin isoforms, HEK293T cells were transfected with constructs expressing HA-tagged tetherin that was visualised by immunofluorescence. *P. alecto* tetherin isoform A localised to the plasma membrane, displaying a similar cellular localisation pattern as human tetherin (Figure 11A and 11C), while isoform C was predominantly localised in the cytoplasm (Figure 11E). *P. alecto* tetherin isoform B demonstrated a fluorescence pattern with features shared by both isoforms A and C (Figure 11D). These data show that bat tetherin isoforms can be expressed with isoforms A and B, but not C predominately localising at the plasma membrane.

**Figure 11.**
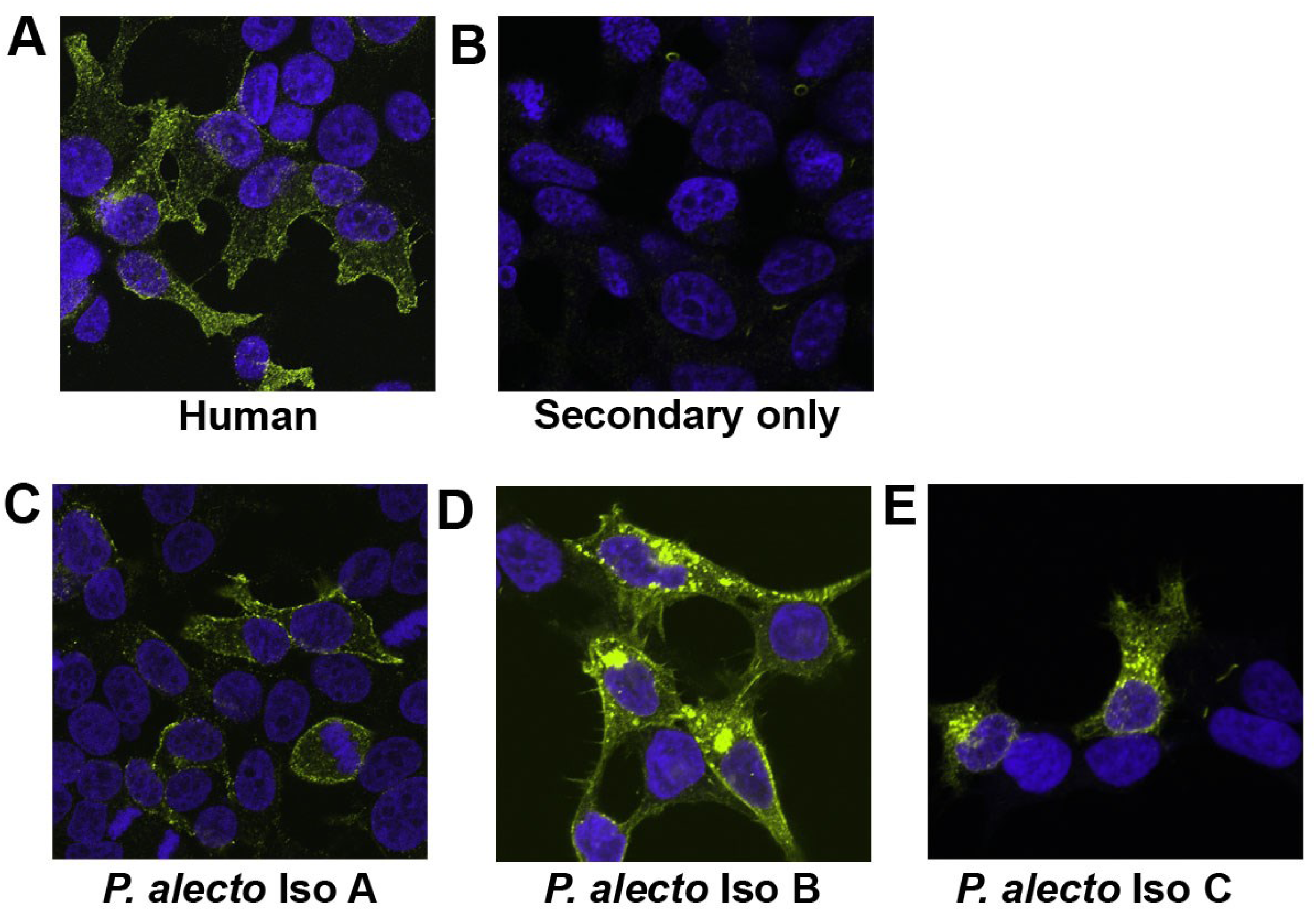
Expression of HA-tagged tetherin proteins. Human and *Pteropus alecto* tetherin isoforms A, B, and C in HEK293T cells. Tetherin localisation in cells was detected using anti-HA-tag rabbit monoclonal Ig and anti-rabbit AlexaFlour 488 secondary Ig. Nuclei are blue stained with Hoechst. Fixed and permeabilised cells were imaged on Nikon AR1 confocal microscope. **A.** Human l-tetherin, **B.** Negative control; HEK293T cells treated without the inclusion of the primary antibody, **C.** *P. alecto* tetherin isoform A, **D.** *P. alecto* tetherin isoform B, **E.** *P. alecto* tetherin isoform C. HA, haemagglutinin.

### *P. alecto* and *M. macropus* tetherin proteins display distinct ability to restrict HIVΔVpu virus-like particles

Tetherin from humans and other mammals including cats, sheep, and other bat species (*Hypsignathus monstrosus* and *Epomops buettikoferi*) restrict the release of HIV-1 VLPs from cells (23, 52, 53). To examine if tetherin from *P. alecto* and *M. macropus* function to similarly block viral particle release, we assessed their ability to restrict HIV-1 VLPs. HEK293T cells were co-transfected with constructs expressing bat tetherin and HIVΔVpu VLPs, that do not express Vpu, an antagonist of human tetherin (21). *P. alecto* tetherin isoforms A and B inhibited the release of HIVΔVpu VLPs in contrast to tetherin isoform C which failed to block VLP release (Figure 12A). *M. macropus* tetherin A also restricted HIVΔVpu VLP release from HEK293T cells; however, tetherin B was unable to inhibit HIVΔVpu VLP egress (Figure 12B). These data indicate that, except for *P. alecto* tetherin isoform C and *M. macropus* tetherin B, HA-tagged bat tetherin proteins are functionally capable of restricting the release of HIV-1 particles from mammalian cells in the absence of Vpu.

**Figure 12.**
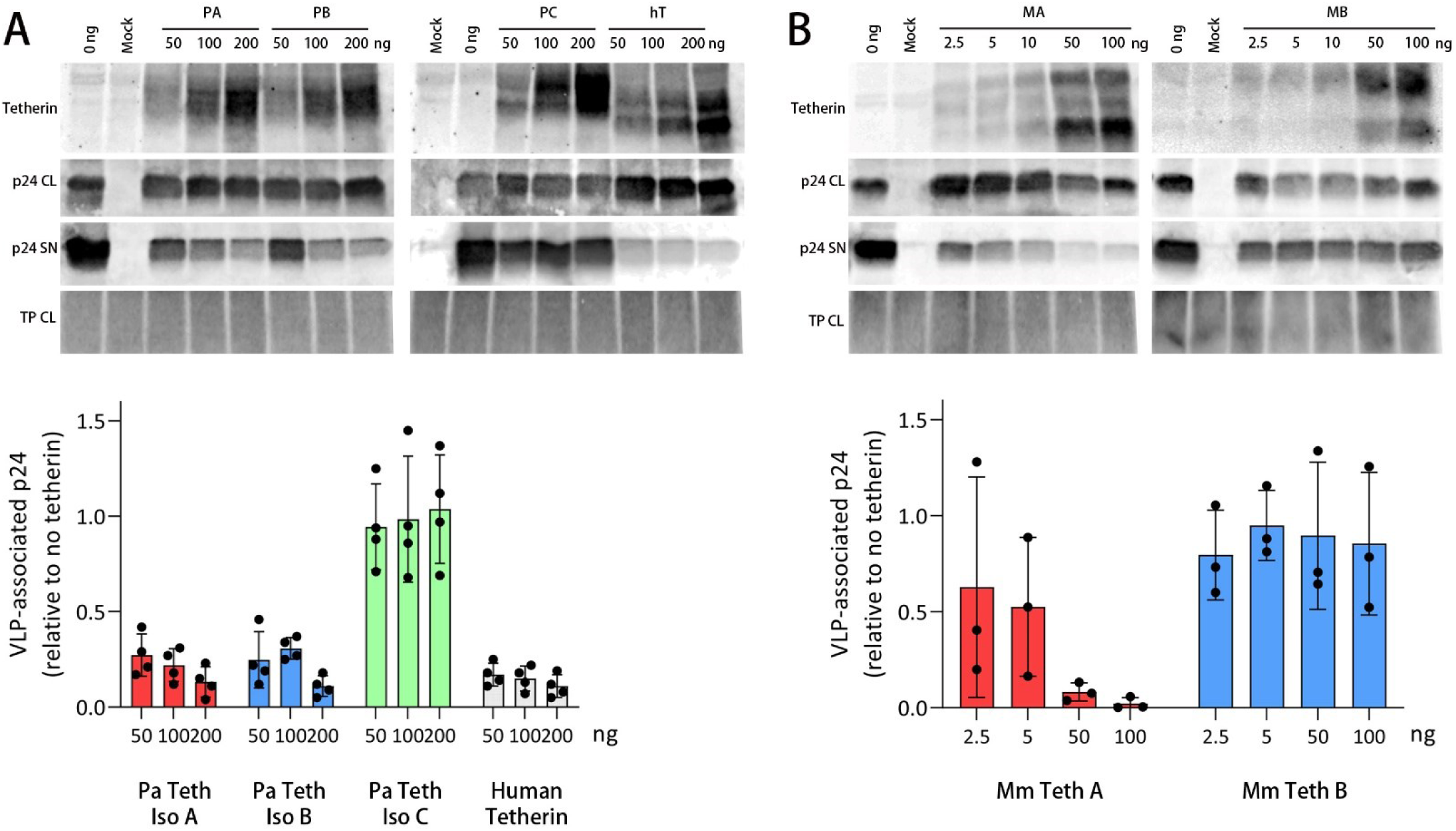
Bat tetherin restricts the release of HIV-1 virus-like particles (VLPs). **A.** *P. alecto* tetherin isoforms A (PA), B (PB), C (PC), or human tetherin, cotransfected with a HIVΔVpu construct. **B.** *M. macropus* tetherin A (MA) and tetherin B (MB) cotransfected with a HIVΔVpu construct. Mammalian HEK293T cells were cotransfected with 200 ng of the HIVΔVpu plasmid expression construct encoding the HIV Gag-Pol polyprotein, which generates HIV-1 VLPs that do not include the tetherin antagonist Vpu, and 0 – 200 ng of the tetherin plasmid expression vector. VLPs were harvested at 48 h and concentrated by ultracentrifugation using a sucrose cushion. VLP and cell lysates were subjected to SDS-PAGE and Western blot analysis. HIV-1 VLPs were detected with a mouse anti-p24 primary antibody and goat anti-mouse Alexa-Fluor 680 fluorophore-conjugated fluorescent secondary antibody. Tetherin was detected with a rabbit anti-HA primary antibody and goat anti-rabbit Alexa-Fluor 800 fluorophore-conjugated fluorescent secondary antibody. Representative Western blots are shown. Total protein stain was used as a loading control. The extent of VLP restriction was quantitated by densitometric analysis of Western blots comparing the relative ratios of VLPs present in the viral lysates and cell lysates from N=4 (*P. alecto*) or N=3 (*M. macropus*) independent assays. Error bars represent the standard deviation. HIV-1, human immunodeficiency virus type 1; SN, cell culture supernatant; CL, cell culture lysate; TP, total protein stain.

### The structurally unique *P. alecto* tetherin isoform C restricts the release of filoviral virus-like particles

We next investigated whether *P. alecto* tetherin isomers are able to restrict filovirus VLPs, including isoform C, which contains a unique C-terminal domain compared to isomers A and B (Figure 3B). In contrast to experiments with HIV-1 VLP, tetherin isoform C was able to restrict the release of VLPs composed of Ebola and Marburg virus VP40 matrix proteins (P = 0.008 for both; Figure 13A & 13B). Ebola VLPs were restricted by isoforms A, B, and C to similar extents (Figure 13A), while Marburg VLPs were restricted by isoform C to a lesser extent than isoforms A and B (Figure 13B). These data demonstrate that all *P. alecto* isomers were able to restrict filoviral VLPs including isomer C which lacks the GSP domain.

**Figure 13.**
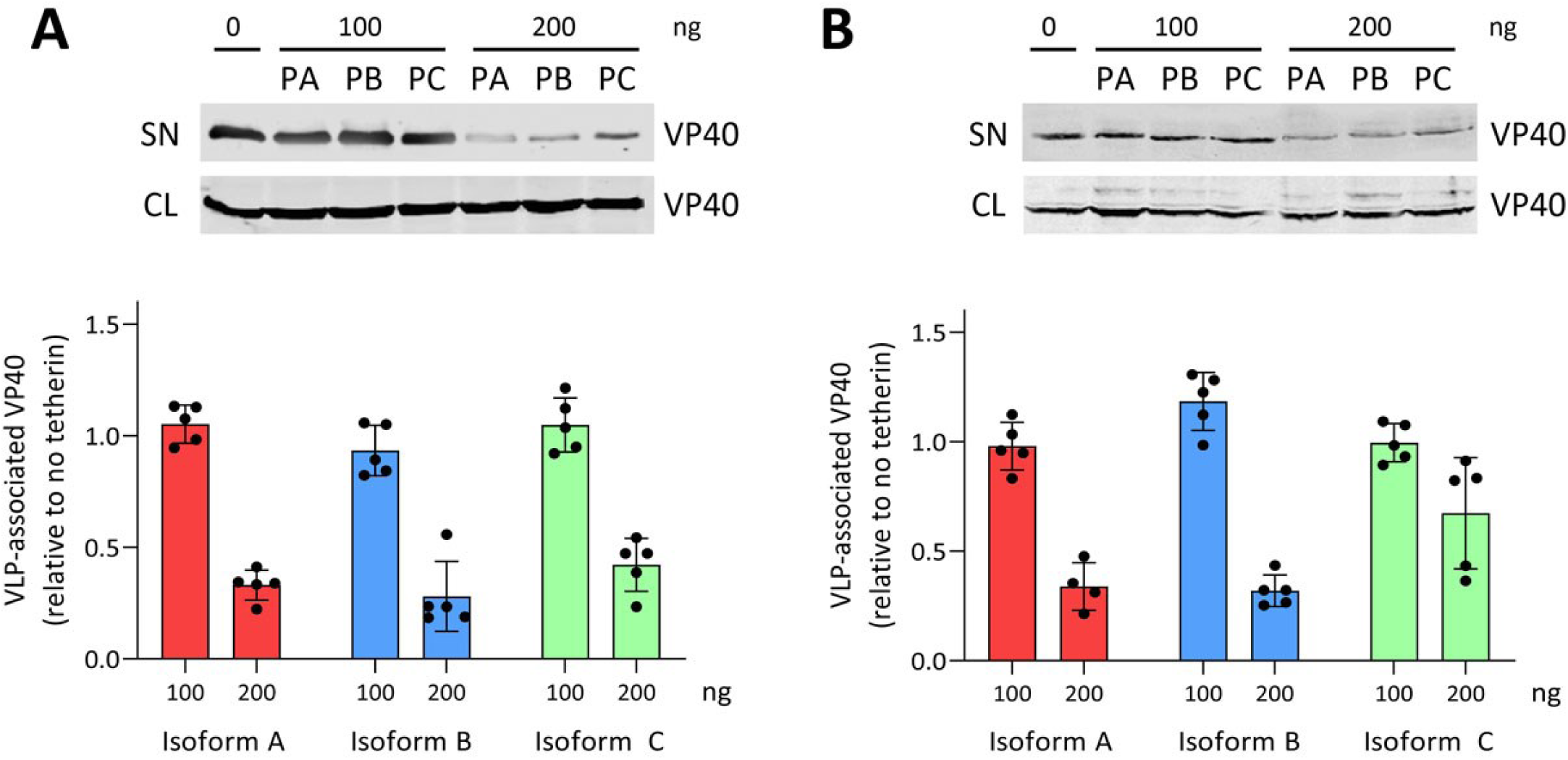
*Pteropus alecto* tetherin restricts the release of Ebola and Marburg virus-like particles (VLPs). *P. alecto* tetherin isoforms A (PA), B (PB), and C (PC), cotransfected with an **A.** Ebola virus construct or **B.** Marburg virus construct. Mammalian HEK293T cells were cotransfected with 200 ng of Ebola or Marburg virus plasmid expression construct encoding a VP40-eGFP protein which generates VLPs, and 0 – 200 ng of the tetherin plasmid expression vector. VLPs were harvested at 48 h and concentrated by ultracentrifugation through a sucrose gradient. VLP and cell lysates were subjected to SDS-PAGE and Western blot analysis. VLPs were detected with a mouse anti-GFP 4B10 primary antibody and a goat anti-mouse Alexa-Fluor 680 fluorophore-conjugated fluorescent secondary antibody. Representative Western blots are shown. The extent of VLP restriction was quantitated by densitometric analysis of Western blots comparing the relative ratios of VLPs present in the viral lysates and cell lysates from N=5 independent assays. Error bars represent the standard deviation. The non-parametric Wilcoxon Rank Sum test was performed to calculate the statistical significance of the restriction of Ebola and Marburg VLPs in the 200 ng tetherin isoform A, B, and C treatment groups; p < 0.01 in all cases. SN, cell culture supernatant; CL, cell culture lysate.

## Discussion

Bats are increasingly being recognised as hosts of viruses with zoonotic potential, driving efforts to better understand these host-virus relationships and the evolutionary features of bats that differentiate them from other mammals (14, 17, 54, 55). It has been hypothesised that the evolutionary adaptation to flight, including changes to the DNA damage response, increased metabolic rates, and higher body temperatures, have influenced the immune system of bats in such a way as to make them ideal hosts for viruses (12, 56, 57). To determine the differences between the innate antiviral defences of bats relative to other mammals, we analysed the genes and expressed transcripts of tetherin from diverse bat genera and species within the order Chiroptera. We found that in all but one species, bats possess genes that express transcripts encoding a tetherin protein homologous to the long isoform of human tetherin (l-tetherin). In addition, we found that *P. alecto* expresses three isoforms from a single tetherin gene, and that vesper bats (genus *Myotis*) encode five, and possibly as many as seven, distinct tetherin genes.

What we have found expands on the findings of Hölzer *et al.,* (18). The three tetherin paralog transcripts identified in *M. daubentonii* correspond to the four paralog genes identified in *M. lucifugus*. The findings of Hölzer *et al.* match our own, as the four genes identified by their group are the tetherin gene regions 1, 2, 6, and 7 described in this study. Region 2 and 6 represent the two different structures observed among *M. macropus* tetherin (Figure 5). Our study expands on these findings in that we describe the expression of five paralogs in *M. macropus* and the identification of seven potential tetherin genes within the tetherin gene locus of *M. lucifugus*. Additionally, we have identified important structural differences between the *Myotis* tetherin paralogs.

The one bat species that lacked the human l-tetherin homolog, *T. melanopogon*, possessed an equivalent of the short isoform of human tetherin (s-tetherin), an observation also reported for cats (58). In a study of feline tetherin, short isoform-exclusivity was found to improve viral restriction and decrease sensitivity to tetherin antagonism by HIV-1 Vpu when compared to an engineered long isoform version of feline tetherin (58). Conversely, the feline immunodeficiency virus (FIV), which is adapted to the feline host, is not restricted by either long or short isoforms of feline tetherin (58). In humans, l-tetherin acts as a virus sensor which induces NF-κB signalling in contrast to s-tetherin, which lacks the dual tyrosine motif required for eliciting this innate immune response (30, 31). The findings presented here suggest that if bat tetherin proteins are confirmed to similarly promote cytoplasmic domain-mediated immune signalling, then *T. melanopogon*, which only encodes a homolog of s-tetherin, would lack this function.

Bat tetherin amino acid sequences were found to be highly variable. The predicted protein lengths range from 177 to 221 AA (human l-tetherin is 180 AA) and of these, only 23 amino acid residues were conserved in 100% of the 27 bat tetherin sequences analysed (Figure 1B). Among these are the structurally important cysteine and asparagine residues which are responsible for tetherin dimerisation and glycosylation, respectively (59, 60). The dual-tyrosine motif, responsible for mediating viral particle endocytosis and immune signalling (31, 40), was found to exist as a variable Y|C·x·Y|H motif across the bat tetherin variants analysed. All bats maintained at least one of the two tyrosine residues. This observation is significant because mutational studies of human tetherin have demonstrated that the dual tyrosines provide redundancy for both the endocytic and signalling activities, which are maintained as long as either tyrosine is present (31, 40).

To understand the evolutionary selective pressures on tetherin genes across bat species, a selection test was performed that revealed that bat tetherin genes are under strong positive selection with amino acid positions in and around the transmembrane domain being the region under the strongest selective pressure (Table 2 and Figure 1A). These data are similar to previous reports that primate tetherin possess multiple sites under positive selection in this same region (28). The driver of positive selection in the case of primate tetherin genes is antagonism by viral counter-measures. These include HIV-1 Vpu (61) and SIV Nef (62), although it is likely that the anti-HIV-1 role of Vpu evolved too recently to have influenced the evolution of human tetherin (62). As virus-host pairs and their respective restriction factors and counter-measures can differ between species within a clade of mammals, such as is the case among primates (61, 62), sites under positive selection in one species may not be under the same pressure in other species. Reflecting this, one analysis of primate tetherin found that it was subject to neutral evolution overall, with episodic adaptive evolution within discrete primate lineages (63). Until more is understood regarding the specific interactions between the tetherin genes of various bat species and the viruses that infect them, we cannot be certain of the drivers of positive selection for bat tetherin. However, it is reasonable to hypothesize that, as for primates, viral antagonists of bat tetherin are likely to exert a large influence.

Venkatesh et al. (64) demonstrated that the configuration that tetherin dimers adopt during viral particle retention primarily consists of the GPI-anchor being embedded in the viral envelope and the transmembrane domain remaining attached to the cellular membrane. If this paradigm holds true for bat tetherin, then it follows that the tetherin-cell membrane interface is the major site of tetherin antagonism in bats and it would be reasonable to speculate that the drivers of this selection are viral antagonists analogous in the mode of interaction, if not structure or function, to lentiviral tetherin antagonists.

We amplified tetherin from spleen-derived cDNA of an Australian fruit bat, *P. alecto*, and confirmed the expression of the two computationally predicted splice variants (isoforms A [X1] and B [X2]), and additionally identified the expression of a third isoform of tetherin, isoform C. Mapped against the *P. alecto* genome, all three isoforms were derived from the alternative splicing of a single tetherin gene (Figure 3B). *P. alecto* tetherin isoform B possesses a 7 AA exclusion within the extracellular coiled-coil domain relative to isoform A, while isoform C is predicted to harbour the same 7 AA exclusion and an alternative C-terminus, that lacks the GSP domain predicted to be present in tetherin isoforms A and B which is necessary for the post-translational addition of the GPI-anchor. This is important because studies of human tetherin have demonstrated that the presence of a GPI-anchor is essential for restricting the release of viral particles (28, 43). The protein encoded by the gene *PLVAP*, which is located downstream of tetherin in *P. alecto* (Figure 4), is tetherin-like in that it encodes a protein with the same structural features as tetherin with the exception of a GSP (29). The human PLVAP protein does not possess restrictive activity against HIV-1 VLPs, but gains restrictive ability when modified to include a GPI anchor (29). Sheep and cows possess a duplication of the tetherin gene (41, 53). In sheep, the duplicate tetherin, named tetherin B, similarly does not encode a GSP, and studies of sheep tetherin B function reveal that while it is capable of limited restriction of VLPs, it is significantly less potent than sheep tetherin A (53). The mechanism of its function is unknown and its C-terminal amino acid sequence is entirely dissimilar from that of *P. alecto* tetherin isoform C. Interestingly, it has been demonstrated that substitution of the GSP with a transmembrane region in human tetherin produces a chimeric protein that retains the ability to inhibit viral particle release (65). However, the C-terminal region of *P. alecto* isoform C is not computationally predicted to include a transmembrane region as determined using a transmembrane helix hidden Markov model, TMHMM 2.0 (66). One possible function of additional isoforms of tetherin is to expand the viral target range of bats by undefined mechanisms that are active in the absence of a GPI anchor.

Intriguingly, one of the non-transmembrane domain-associated sites under strong positive selection across tetherin from bats is located in the middle of the extracellular domain. This is the site that distinguishes the *P. alecto* tetherin isoforms A and B, with the 7 AA exclusion at this location in isoform B. This suggests that the expression of *P. alecto* tetherin isoform B, and possibly also *M. macropus* tetherin B and E, which differ from tetherin A through a large 60 AA deletion in the same region, is to express a tetherin variant lacking sequence motif that appears to be a possible site of viral antagonism. This hypothesis could be supported by future experiments aimed at identifying viral antagonists of *P. alecto* tetherin isoform A that are ineffective against isoform B.

We found that the microbat *M. macropus* expresses five different tetherin genes, tetherin A, B, C, D and E, of which three (tetherin A, C, and D), encode homologs of human l-tetherin, while *M. macropus* tetherin B and E possess a large 60 AA deletion within the extracellular coiled-coil domain (Figure 4). Mapping of these genes against the genome assembly of the relatively distantly related *M. lucifugus* indicates that vesper bats of the genus *Myotis* may possess as many as seven tetherin genes (Figure 6). The presence of multiple tetherin genes suggests that these bats have expanded and diversified their use of these antiviral restriction factors relative to other mammals, supporting the hypothesis of differential antiviral gene evolution in bats.

Tetherin was expressed widely and variably across the tissues of *P. alecto*, which is consistent with previous observations of human tetherin (36). We observed the highest levels of *P. alecto* tetherin expression in the thymus, possibly reflecting the role of the thymus in expressing a large proportion of the proteome due to its role in central tolerance. High levels of expression were also observed in lung tissue, particularly in the lung of an individual female bat. This may reflect a frontline defensive role of tetherin in the lung against enveloped viruses that are respiratory pathogens.

Tetherin was upregulated in *P. alecto* spleen tissue treated with TLR agonists LPS and PIC. Interestingly, we observed that the expression of isoforms A, B, and C was highly variable among individual spleen samples. In several spleens, only isoform B was expressed, while in one spleen treated with PIC the vast majority of expression was attributed to isoform A. The remainder expressed variable levels of each isoform A and B, with the majority of expression trending toward isoform B. Surprisingly, isoform C was only expressed, at a low level, in a single spleen sample that had been treated with PIC. The biological importance of this observation is not presently known. Ongoing assessments of bat tetherin expression should determine if this strong bias in alternative isoform expression is present in other tissues, such as the thymus and lung.

The expression of the *P. alecto* tetherin isoforms A, B, and C, and *M. macropus* tetherin A and B, reveals differences in their relative capacities to form homodimers under our assay conditions (Figure 9A). This observation is notable because the ability of tetherin to form cysteine-linked dimers is required for the restriction of HIV-1 viral particles (60) which we chose as the VLPs against which bat tetherin proteins were functionally validated for inhibiting viral particle release. In contrast to restriction of HIV-1, tetherin dimerisation is not required for the restriction of arenaviruses and filoviruses (67), suggesting that the need for dimerisation is virus-dependent.

The *P. alecto* tetherin isoform A, which is homologous to human l-tetherin, was found to predominantly form dimers (Figure 9A), which is consistent with human tetherin (Figure 9B). In contrast, *P. alecto* tetherin isoform B and C, and *M. macropus* A was also observed in monomeric form under non-reducing conditions (Figure 9A). *P. alecto* isoforms B & C contain a 7 AA exclusion in their ectodomain relative to isoform A, and isoform C possesses a distinct C-terminal region, which may account for this observation by potentially impacting dimer formation and/or stability. Additionally, we cannot currently rule out the possibility that the insertion of the HA tag may influence tetherin dimerisation. *M. macropus* tetherin B was observed exclusively as a monomer (Figure 9A). The monomeric exclusivity of *M. macropus* tetherin B can be explained by the loss of all conserved cysteine residues necessary for dimerisation within the 60 AA deletion in the extracellular coiled-coil domain of tetherin B (Figure 4). Bat tetherin proteins were observed to be less sensitive to reduction by dithiothreitol compared to human tetherin (Figure 9C). One possible explanation for this is the predicted presence of a region of hydrophobicity at the N-terminus (position 1-10 AA) of *P. alecto* and *M. macropus* that was not predicted for human tetherin (Figure 10); however, other differences in the secondary structure of bat vs. human tetherin may also account for this effect.

Given the differing extent of homodimer formation for each bat tetherin and reports that dimerisation is important for the restriction of some viruses and not required for others (60, 67), it is likely that these differences might affect the extent by which each tetherin is capable of restricting various VLPs. Tetherin function was validated through an assessment of the capacity of *P. alecto* tetherin isoforms A, B, and C, and *M. macropus* tetherins A and B to inhibit the egress of HIVΔVpu VLPs. While primate lentiviruses are not known to infect bats, bats have recently been discovered to host extant gammaretroviruses and deltaretroviruses, and bat genomes contain diverse endogenous retroviral sequences (27, 68–71). *P. alecto* tetherin isoforms A and B inhibited the release of HIVΔVpu VLPs, in contrast to isoform C which lacked restriction activity (Figure 12A). *M. macropus* tetherin A restricted HIVΔVpu VLP release, while tetherin B did not (Figure 12B). These findings are consistent with the expected effects of tetherin dimerisation on the restriction of HIVΔVpu VLPs (60).

Previous reports on the necessity of the tetherin GPI anchor for inhibition of viral particle release (28, 43) indicate that the lack of a GPI anchor on *P. alecto* isoform C would likely result in an inability to inhibit viral egress from the host cell. However, sheep tetherin B, which does not possess a GPI-signal peptide (Figure 2B), is capable of restricting the release of betaretroviruses by blocking glycoprotein incorporation into nascent virions (53, 72). Because of the unique sequence of the C-terminus and lack of GSP in *P. alecto* tetherin isoform C, we extended our evaluation of its restrictive capacity to include filoviral VLPs derived from Ebola and Marburg virus VP40 matrix proteins. Surprisingly, *P. alecto* tetherin isoform C was capable of inhibiting the egress of Ebola VLPs to an extent similar to isoforms A and B, although it was less restrictive of Marburg VLPs compared to isoforms A and B (Figure 13).

How isoform C could be capable of restricting the release of VLPs without a C-terminal GPI anchor is not presently known, although there are at least two possible explanations. The first is that the alternative C-terminal sequence of isoform C is involved in envelope or cellular membrane binding through an unknown mechanism. The second is that isoform C forms dimers in a manner distinct from that of isoforms A and B. One possibility is that it can form an antiparallel homodimer configuration in such a way that the N-terminus of one monomer aligns with the C-terminus of another monomer, forming a dimer with a transmembrane domain at each end of the protein. Another possibility is the formation of standard parallel dimers of tetherin followed by the formation of an antiparallel tetramer, similarly resulting in a quaternary structure possessing a transmembrane domain at either end. This possibility is supported by structural analyses demonstrating that human tetherin is capable of forming tetramers composed of antiparallel pairs of parallel dimers (59, 73).

## Conclusions

Tetherin variants of the fruit bat *P. alecto* and the vesper bat *M. macropus* are functionally capable of restricting the release of VLPs in a mammalian cell culture system. These bats have evolved to express unique forms of tetherin that do not conform to the standard tetherin protein structure observed in tetherin proteins of other mammals. A structurally unique tetherin of *P. alecto*, isoform C, was not observed to inhibit the egress of HIVΔVpu VLPs but was capable of restricting filoviral VLPs. These findings raise questions regarding the possible antiviral range, mechanism of action, and *in vivo* function of structurally diverse forms of tetherin. Bat tetherin genes are under strong positive selection, indicating an ongoing process of evolution involving tetherin targets and viral antagonists. This evolutionary pressure has resulted in the expansion and diversification of tetherin genes in vesper bats of the genus *Myotis*, and the emergence of unique splice variants of tetherin in the fruit bat *P. alecto*, supporting the hypothesis of differential and unique antiviral gene evolution in bats.

## Methods and Materials

### Sequence read archive (SRA) BLAST analysis

To predict the tetherin sequences of other bat species, the *P. alecto* tetherin isoform A coding domain nucleotide sequence (549 nt) was used as the search query for a separate BLASTn analysis of each of the publicly accessible sequence read archives (SRA) of bat transcriptomes (Table 1). This approach allowed the individual assembly of a tetherin homolog from each bat species for which a transcriptome sequence read archive was available. The BLASTn analyses were performed using the online SRA Nucleotide BLAST tool (https://blast.ncbi.nlm.nih.gov/Blast.cgi). The program selection was optimised for ‘somewhat similar sequences’ (BLASTn) and the selected algorithm parameters were as follows: Max target sequences = 1000; Expect threshold = 1.0x10^-10^; Word size = 7; Max matches in a query range = 21; Match/Mismatch scores = 2,-3; Gap costs of existence 5 and extension 2; and no filtering or masking was selected. Matching reads were downloaded and assembled using the Assemble Sequences tool in CLC Genomics Workbench 8.0 (CLC; Qiagen, Germany, https://www.qiagenbioinformatics.com/). The consensus sequence of the assembly was then used as the search query for a second SRA BLAST analysis with the same parameters as the first search with the following exceptions: Program selection was optimised for ‘highly similar sequences’ (megablast); Expect threshold = 1.0 x 10^-20^; Word size = 28; Match/Mismatch scores = 1,-2; and linear gap costs of existence and extension. Matching reads were downloaded and assembled in the same manner, and the assembled consensus sequence was used in a third SRA BLAST using the same parameters as the second. This process was iteratively repeated for the assembled consensus sequence of each bat tetherin until it extended through the tetherin coding domain in both directions into the 5’ and 3’ untranslated regions, respectively demarcated by the locations of the start methionine and stop codon.

### Evolutionary selection test

To determine if evolutionary selective pressures were being applied to bat tetherin, a selection test was performed. To detect the positively selected sites among bat tetherin sequences, a maximum likelihood (ML) phylogeny was generated with CODEML implemented in PAML4 software (74). The input tree was generated based on phylogeny generated by TimeTree (75). The coding sequence alignments were fit to the NSsites models allowing (M8; positive selection model) or disallowing (M8a; null model) positive selection. Models were compared using a chi-squared test (degrees of freedom = 2) on twice the difference of likelihood values to derive P-values. The codon frequency model F3x4 was used. In cases where a significant difference (P < 0.01) between M8a versus M8 was detected, the Bayes Empirical Bayes (BEB) analysis was used to identify codons with ω (dN/dS) > 1, reporting values with posterior probability > 0.99 as high significance or 0.95 - 0.99 as moderate significance.

### Identification of protein domains, structures, and motifs

To confirm the predicted tetherin nucleotide sequences and identify functional domains within bat tetherin protein sequences, translated coding domains were compared against known tetherin amino acid sequences including the human l-tetherin and the *P. alecto* tetherin isoform A (Figure 2). For all tetherin amino acid sequences, secondary structures were determined using the Predict Secondary Structure tool in CLC. The presence of cytoplasmic, transmembrane, and extracellular domains were determined using a hidden Markov model in TMHMM 2.0 (66). GPI signal peptides (GSP) were determined using the online tool PredGPI (76) (http://gpcr2.biocomp.unibo.it/gpipe/index.htm). Multiple sequence alignments (MSA) of predicted and known tetherin amino acid and nucleotide sequences were performed using MUSCLE v3.8 (77) with the following parameters: Maximum iterations = 16, Find diagonals = No. The molecular weight of unglycosylated tetherin proteins with the GSP removed were calculated using the Expasy Compute pI/Mw online tool (78) (https://web.expasy.org/compute_pi/) with the addition of 1.5 kDa to account for the presence of the GPI anchor (79). Glycosylation sites were predicted using the GlycoMine online tool (50) (https://glycomine.erc.monash.edu/Lab/GlycoMine/). Tetherin protein hydrophobicity was analysed using the hydrophobicity plot tool in CLC with the following parameters: Applied scale = Engleman; Window size = 11.

### Transcriptome and contig analysis

Approval for the use of bat tissue was granted by the Australian Centre for Disease Preparedness (ACDP) (formerly the Australian Animal Health Laboratory, AAHL) Animal Ethics committee (Protocol AEC1281). The *P. alecto* transcriptome is accessible through the NCBI Sequence Read Archive (http://www.ncbi.nlm.nih.gov/Traces/sra/) [SRA: SRP008674].

To identify the homologs of tetherin in *P. alecto*, the human l-tetherin protein sequence [UniProt: Q10589-1] was used as a query in a tBLASTn analysis of the transcriptome of *P. alecto*. To identify assembled sequences (contigs) representing mRNA transcripts of interest, local tBLASTn analyses of the transcriptome of *P. alecto* were conducted with CLC using the following parameters: BLOSUM62 matrix, word size = 3, E-values < 1x10^-12^, gap costs of existence 11, extension 1, with no filtering of regions of low complexity.

### cDNA analysis

To amplify tetherin nucleotide sequences from bat cDNA, polymerase chain reaction (PCR) assays were performed using cDNA generated from *P. alecto* and *M. ricketti* spleen tissue, and *M. macropus* and *M. schreibersii* kidney cell lines, using various combinations of forward and reverse primers (Table 3A). Primers were designed using the tetherin predictions identified in the contig and SRA analyses. Primers were designed to bind to the 5’ and 3’ untranslated regions of bat cDNA such that the full CDS could be amplified. Bat capture, tissue collection and RNA extraction was conducted as previously reported (80) with the exception that RNAlater (Ambion, USA) preserved spleen tissue from four male adult bats was pooled before tissue homogenisation and followed with total RNA extraction with the Qiagen RNeasy Mini kit with on-column digestion of genomic DNA with DNase I. Total RNA was reverse transcribed into cDNA with the Qiagen Omniscript reverse transcriptase according to the manufacturer’s protocol with the exception that the reaction contained 100 ng/μl total RNA, 1 μM oligo-dT18 (Qiagen) and 10 μM random hexamers (Promega).

All PCR amplification assays were performed using the Roche FastStart High Fidelity PCR system (Cat # 04738292001) with an annealing temperature gradient of 54°C to 64°C in 2°C increments. Each reaction was made up to a total of 20 μl, containing 1 unit of polymerase, 2 ng of total cDNA, and 8 pmol of each primer. All other parameters for PCR amplification were performed according to the manufacturer’s protocol.

Amplicons from PCR reactions were analysed by agarose gel electrophoresis (81). Individual DNA bands were physically excised from the gel. DNA was purified from the gel fragments using the Wizard SV Gel and PCR Clean Up kit (Promega, Fitchburg, USA) according to the manufacturer’s protocol.

To further analyse the PCR amplified DNA, each DNA fragment was blunt-end ligated into the pCR2.1-TOPO-TA or pCR-Blunt-II-TOPO plasmid vector (Invitrogen, Waltham, USA). Ligation was performed using the TOPO TA or Zero Blunt TOPO PCR Cloning Kit (Invitrogen) according to the manufacturer’s instructions and plasmids were transformed into Top10 *E. coli* using a standard heat-shock method (81). Plasmids were purified from these cultures using a Wizard Plus SV Miniprep DNA Purification kit (Promega). All inserted sequences were confirmed by Sanger sequencing using M13 forward and reverse primers (M13F and M13R; Table 3B).

### Genome Mapping

To identify tetherin genes within the genomes of *P. alecto* and *M. macropus*, local BLASTn analyses were performed using CLC. For these analyses the *P. alecto* and *M. macropus* tetherin sequences were used as query sequences against the *P. alecto* genome [NCBI: PRJNA171993] and *M. lucifugus* genome [NCBI: GCF_000147115], respectively. BLASTn was performed using the following parameters: Word size = 11, E-values < 1x10^-3^, gap costs of existence 5, extension 2, with no filtering of regions of low complexity. Genes were delineated as beginning and ending at the first and last nucleotide of the contig query sequence. Exons were delineated as consisting of the nucleotide regions within the gene mapping to the tetherin cDNA sequences, bordered by the canonical 5’-AG and 3’-GT dinucleotide motifs (82). The 5’ and 3’ untranslated regions were defined as the regions upstream of the start methionine and downstream of the stop codon of the CDS, respectively. Contig sequences were mapped against gene scaffolds through a local BLASTn analysis using CLC Genomics Workbench with default settings. The nucleotide sequence preceding the first exon of each gene was assessed for the presence of promoter motifs.

Neighbouring genes were identified by BLAT (BLAST-like alignment tool) analysis (83) of the bordering nucleotide sequences upstream and downstream of tetherin genes using the Ensembl genome database BLAT tool (http://asia.ensembl.org/Multi/Tools/Blast?db=core).

### Generation of tagged tetherin constructs for expression in mammalian cells

To enable detection of tetherin protein expression, the *P. alecto* tetherin isoforms A, B, and C, and *M. macropus* tetherin A and B were genetically modified with the insertion of nucleotide sequences encoding the haemagglutinin (HA) antibody-binding epitope. Tetherin sequences were modified through a 2-step PCR process. PCR reactions were performed using the Roche FastStart HighFidelity PCR kit according to the manufacturer’s recommendations.

To express tetherin proteins in a mammalian cell culture system the tagged tetherin inserts were sub-cloned from the pCR2.1-TOPO (*P. alecto* tetherin isoforms A, B, and C) and pCR-Blunt-II-TOPO (*M. macropus* tetherin A and B) vectors into pcDNA3.1 mammalian expression vectors. The HA-tagged tetherin constructs contained terminal enzyme restriction sites. *P. alecto* tetherin constructs contained *Xho*I and *Xba*I sites at their 5’ and 3’ ends, respectively. *M. macropus* tetherin constructs contained an *Eco*RI site at each end. These sites, which are also present in the pcDNA3.1 vector, were used for digestion-ligation transfer of the HA-tagged tetherin sequences into pcDNA3.1. The ligation reaction products were transformed into Top10 *E. coli* and plasmid clones were purified from *E. coli* colonies using the method described under ‘cDNA analysis’. All inserted sequences were verified by Sanger sequencing using the pcDNA3.1 vector sequencing primers, T7F forward and BGHR reverse (Table 3B).

### Expression, extraction, and detection of tetherin in a mammalian cell culture system

To determine if tetherin plasmids express tetherin protein in a mammalian cell culture system, adherent human embryonic kidney HEK293T cells (kindly provided by Richard Axel, Columbia University) were transfected with each tetherin construct with protein expression determined by Western blot analysis.

HEK293T cells were maintained in Dulbecco’s modified Eagle medium (DMEM-10, Thermo Fisher) enriched with heat-inactivated foetal calf serum (100ml/L; Invitrogen), glutamine (292 μg/ml; Invitrogen), and the antibiotics penicillin (100 units/ml; Invitrogen) and streptomycin (100 µg/ml; Invitrogen) (DMEM-10). Cells were incubated at 37°C with 5% CO2.

The expression was performed in 6-well plates. Each well was seeded with 3 x 10^5^ cells/well in 2 ml of DMEM-10. Cells were transfected when the monolayer had reached 50-60% confluency. The tetherin constructs analysed are listed in Table 4, except for the human tetherin construct, pTethHA463, which has been previously described (61). Cells were transfected with each plasmid in duplicate wells at 2 μg/well using Lipofectamine 2000 (Thermo Fisher) according to the manufacturer’s protocol. Tetherin was extracted using a previously published GPI-anchored protein extraction protocol (49).

Protein samples were analysed by size-based separation through SDS-PAGE using either 12% polyacrylamide gels or Any kD polyacrylamide gels (Bio-Rad) (81). Samples run under reducing and deglycosylating conditions were reduced by treatment of the samples with dithiothreitol (final concentration, 100 mM) and deglycosylated with PNGase F (New England Biolabs, Ipswich, USA), respectively following the manufacturer’s instructions. Proteins were transferred to Amersham Protran nitrocellulose membranes (Sigma) for Western blot analysis. Revert Total Protein stain (LI-COR Biosciences, Lincoln, USA) was applied to the membranes, and visualised using the Odyssey Imaging System (LI-COR) according to the manufacturer’s protocol. For the Western blot analysis, the primary antibody solution contained a 1/1000 dilution of a monoclonal rabbit anti-HA antibody (C29F4, Cell Signaling Technology, Danvers, USA) in Tris-buffered saline (TBS; pH 7.6) containing 0.1% Tween-20, and the secondary antibody solution contained a 1/10,000 dilution of a polyclonal goat anti-rabbit IRD800 fluorophore-conjugate secondary antibody (Rockland, USA). The primary antibody solution was incubated overnight at 4°C, and the secondary antibody solution was incubated at room temperature for 1 h. To visualise fluorescent antibody-bound tetherin proteins, membranes were scanned using the Odyssey Imaging System (LI-COR) at wavelengths of 600 and 800 nm, using the default software settings.

### Fluorescence microscopy

To visualise tetherin localisation within cells, 500 ng of plasmids encoding HA-tagged human l-tetherin and *P. alecto* tetherin isoforms A, B, and C were transfected into HEK293T cells seeded on glass coverslips as described above. At 48 h post transfection, cells were fixed with 4% paraformaldehyde (Sigma) in PBS for 10 min at room temperature, and then permeabilised in 0.2% Triton X-100 in PBS for 5 min at room temperature. Tetherin localisation in cells was detected by staining cells with anti-HA-tag rabbit monoclonal IgG (Thermo Fisher) diluted in 0.2% Triton X-100, 3% bovine serum albumin (BSA) in PBS for 1 h at room temperature. Cells were subsequently stained with anti-rabbit AlexaFlour 488 secondary IgG (Thermo Fisher) diluted in 0.2% Triton X-100, 3% BSA in PBS for 20 min at room temperature. Nuclei were blue stained with Hoechst 33342 (Thermo Fisher) according to the manufacturer’s instructions. Coverslips were mounted onto glass slides with ProLong Gold Antifade Mountant (Thermo Fisher), then imaged on a Nikon AR1 confocal microscope and analysed with ImageJ (NIH) software.

### qPCR analysis of tetherin expression across multiple bat tissues

To determine tetherin expression across various *P. alecto* tissues, tetherin mRNA levels were measured by qPCR analysis. The primers and probes were designed using the program Primer Express (Perkin–Elmer, Applied Biosystems, USA). The tetherin primers amplify all three known isoforms of *P. alecto* tetherin. *P. alecto* bats were trapped in Queensland, Australia, and transported alive by air to the ACDP in Victoria, where they were euthanised for dissection using methods approved by the ACDP animal ethics committee (AEC1389). Tissues were stored at −80 ◦C in RNAlater (Ambion). Total RNA was extracted from frozen *P. alecto* tissues using a Precellys 24 tissue homogeniser (Bertin Technologies, France) and a RNeasy mini kit (Qiagen) with on-column DNase-I treatment (Qiagen) to remove traces of genomic DNA.

Total RNA was subjected to real-time PCR using SuperScript III One-Step RT-PCR System (Invitrogen) with Platinum *Taq* DNA Polymerase. Reactions were performed on 100 ng of template RNA with 200 nM of each primer and 150 mM of the Taqman probe in an Applied Biosystems 7500 Fast Real-Time qPCR instrument. Cycling parameters consisted of 50°C, 5 min for the reverse transcription of RNA to cDNA followed by 95°C for 2 min. The cDNA was amplified by PCR for 40 cycles, each consisting of 95°C for 3 s and 60°C, 30 s. Using the Livak/ΔΔ Ct method (84), tetherin Ct values were normalised against the expression/Ct values of the 18S rRNA housekeeping gene and reported as fold-difference, calibrated against the Ct values for wing tissue.

### Transcriptome analysis of isoform expression under immune-stimulating treatments

To compare the expression of alternative isoforms of *P. alecto* tetherin and the impact of treatment with immune-stimulating compounds, bats were treated with PBS, LPS (Invivogen, San Diego, USA) or PIC (Invivogen) as published previously (47). Briefly, 5 h post-intraperitoneal injection, bats were anaesthetised, culled and organs were processed for RNA, DNA, protein and cell suspensions as described previously (85). RNA libraries were prepared using RiboZero Plus rRNA-depletion kits (Illumina, USA) and cDNA was generated using a mix of oligo-dT/random hexamer primers, prior to sequencing in 2x150PE on the Illumina HiSeq platform.

Sequencing read libraries were quality controlled by analysis using FastQC (86). Illumina sequence adapters were removed and reads were trimmed or discarded on the basis of quality using the Trim Sequences tool in CLC. Overlapping paired reads were merged using the Merge Overlapping Pairs tool in CLC. Using the RNA-Seq Analysis tool in CLC, sequence reads were mapped against the *P. alecto* gene scaffold containing the tetherin gene (GenBank accession: KB030270.1), which was manually annotated with the tetherin gene and mRNA sequences for isoforms A, B, and C. The following parameters were used for the RNA-Seq analysis: mismatch = 3, insertion = 3, deletion = 3, length fraction = 0.6, similarity fraction = 0.95; default parameters were used otherwise. Isoform expression levels were normalised for post-trimming read library size and reported as counts-per-million reads (CPM). Non-parametric one-tailed Mann-Whitney tests were performed to calculate statistical significance between treatments. The *P. alecto* tetherin gene scaffold, annotation files, and read maps are provided in SI Data 2.

### Functional validation of tetherin activity

To determine the inhibitory effect of bat tetherin on the release of virus-like particles (VLPs), HEK293T cells were co-transfected with each tagged bat or human tetherin construct and either a HIVΔVpu plasmid construct, pCRV1-NLgagpolΔVpu, which expresses HIV NL4.3 Gag-polΔVpu protein [NCBI: P12497.4]; or an EBOV VP40 plasmid, pGFPEVP40, expressing the Ebola Zaire virus VP40 Matrix protein [NCBI: NP_066245]; or a MARV VP40 plasmid, pGFPEMVP40, expressing the Marburg virus VP40 Matrix protein [NCBI: YP_001531155]. EBOV and MARV VP40 proteins were encoded and expressed as VP40-eGFP fusion proteins. VLP plasmid and human tetherin constructs were kindly provided by Paul Bieniasz (Aaron Diamond AIDS Research Center, Rockefeller University) (87).

Transfections were performed as described above, with the following modifications: 12-well plates were seeded with 1.5 x 10^5^ cells/well in a total volume of 1 ml of DMEM-10, DNA-Lipofectamine mixtures for each well contained 1 μl of Lipofectamine, and a total of 400 ng of DNA comprised of 200 ng of VLP plasmid DNA, and 0 – 200 ng of tetherin-encoding plasmid DNA, made up to a total mass of 200 ng with insert-free pcDNA3.1 plasmid DNA. Each sample was prepared in duplicate wells. Cells were incubated at 37°C with 5% CO2 for a total of 48 h. After 24 h, 1 ml of DMEM-10 was added to each well. Following incubation cell lysates and supernatants were collected for further analysis.

Cell culture supernatants were collected and clarified. Clarified supernatants were layered over 2 ml of a 25% (w/v) sucrose solution in 13.2 ml Thinwall Polypropylene SW41 Ti ultracentrifuge tubes (Beckman-Coulter, Brea, USA) and made up to a total volume of 10 ml with PBS without calcium chloride or magnesium chloride (PBS[-]). Samples were then centrifuged at 130,000 x *g* for 2 h at 4°C in the Optima L-100 XP ultracentrifuge (Beckmann-Coulter). VLP pellets were lysed in 60 μl of NP-40 lysis buffer containing 1% NP-40 (Sigma-Aldrich) and 10 μg/ml each of aprotinin, leupeptin, and pepstatin A protease inhibitors (Sigma-Aldrich).

The cells from each well were resuspended in 500 μl of PBS[-] and transferred to 1.5 ml tubes and centrifuged at 200 x *g* for 5 min. Cell pellets were lysed in 100 μl of NP-40 lysis buffer. Following cell lysis samples were clarified by centrifugation at 20,000 x *g* for 5 min at 4°C, and supernatants collected for SDS-PAGE.

Cell and viral lysates were analysed by SDS-PAGE and Western blot analysis as described above, with the following exceptions. For HIV VLPs, the primary antibody binding was directed against the Gag protein using a solution containing a 1/4000 dilution of a mouse anti-p24 antibody (NIH AIDS Reagent Repository, Germantown, USA) and 0.1% Tween-20 in PBS[-]. For Ebola and Marburg VLPs, the primary antibody binding was directed against eGFP using a solution containing a 1/4000 dilution of a mouse anti-GFP 4B10 antibody (Cell Signaling Technology, USA). For all VLPs, the secondary antibody was a 1/10,000 diluted polyclonal goat anti-mouse Alexa-Fluor 680 fluorophore-conjugate fluorescent secondary antibody (Thermo Fisher).

VLPs released into the supernatant for each sample were quantified using a densitometric analysis of the signal strength of the fluorescent antibodies bound to VLP proteins. Densitometry was performed using the Image Studio Lite v5.2 software suite (LI-COR). Variation in viral lysate signal strength across samples was normalised using the signal strength of VLP proteins in the cell lysates. The non-parametric Wilcoxon Rank Sum test was performed to calculate the statistical significance of the restriction of Ebola and Marburg VLPs in the 200 ng tetherin isoform C treatment groups.

### Data availability

The data underlying this article are available in this article and its online supplementary material. Novel tetherin sequences have been deposited in GenBank, at https://www.ncbi.nlm.nih.gov/genbank/ with accession numbers listed in Table 1.

## Acknowledgments

We thank Paul Bieniasz for providing the pCRV1-NLgagpolΔVpu and pTethHA463 plasmid constructs. We thank Richard Axel for providing HEK293T cells. This work was supported by the National Health and Medical Research Council (NHMRC), grant number GNT1121077, awarded to G.T., M.L.B., and G.A.M., and NHMRC Senior Research Fellowship GNT1117748 to G.T. J.A.H. was funded by NHMRC GNT1121077 and a Monash University Vice Chancellor’s Honours-PhD scholarship. L-F.W. is funded by the Singapore National Research Foundation grants NRF2012NRF-CRP001-056 and NRF2016NRF-NSFC002-013. J.C. is funded by National Natural Science Foundation of China (31671324) and CAS Pioneer Hundred Talents Program. The authors gratefully acknowledge the contribution to this work of the Victorian Operational Infrastructure Support Program received by the Burnet Institute.

## Author’s Contributions

J.A.H., M.T., L-F.W., and G.T. designed the study; J.A.H., J.C., A.T.I, A.N., I.S., T.B.G. and A.J., performed research and analysed data; J.A.H., M.T., A.J., A.N., V.B., G.A.M., M.L.B., L-F.W., and G.T. discussed and interpreted the results; J.A.H. wrote the first draft of the paper; J.A.H., and G.T. critically edited the manuscript.

## References

1. Calisher CH, Childs JE, Field HE, Holmes KV, Schountz T. 2006. Bats: Important reservoir hosts of emerging viruses. Clin Microbiol Rev 19:531–545.

2. Smith I, Wang L-F. 2013. Bats and their virome: an important source of emerging viruses capable of infecting humans. Curr Opin Virol 3:84–91.

3. Amman BR, Albariño CG, Bird BH, Nyakarahuka L, Sealy TK, Balinandi S, Schuh AJ, Campbell SM, Ströher U, Jones ME. 2015. A recently discovered pathogenic paramyxovirus, Sosuga virus, is present in *Rousettus aegyptiacus* fruit bats at multiple locations in Uganda. J Wildl Dis 51:774–779.

4. Zhou P, Yang X-L, Wang X-G, Hu B, Zhang L, Zhang W, Si H-R, Zhu Y, Li B, Huang C-L. 2020. A pneumonia outbreak associated with a new coronavirus of probable bat origin. Nature:1–4.

5. Letko M, Seifert SN, Olival KJ, Plowright RK, Munster VJ. 2020. Bat-borne virus diversity, spillover and emergence. Nat Rev Microbiol:1–11.

6. Swanepoel R. 1996. Experimental inoculation of plants and animals with Ebola virus. Emerg Infect Dis 2:321.

7. Williamson MM, Hooper PT, Selleck PW, Gleeson LJ, Daniels PW, Westbury HA, Murray PK. 1998. Transmission studies of Hendra virus (equine morbilli-virus) in fruit bats, horses and cats. Aust Vet J 76:813–818.

8. Middleton DJ, Morrissy CJ, van der Heide BM, Russell GM, Braun MA, Westbury HA, Halpin K, Daniels PW. 2007. Experimental Nipah virus infection in pteropid bats (*Pteropus poliocephalus*). J Comp Pathol 136:266–272.

9. Watanabe S, Masangkay JS, Nagata N, Morikawa S, Mizutani T, Fukushi S, Alviola P, Omatsu T, Ueda N, Iha K, Taniguchi S, Fujii H, Tsuda S, Endoh M, Kato K, Tohya Y, Kyuwa S, Yoshikawa Y, Akashi H. 2010. Bat coronaviruses and experimental infection of bats, the Philippines. Emerg Infect Dis 16:1217–1223.

10. Jones ME, Schuh AJ, Amman BR, Sealy TK, Zaki SR, Nichol ST, Towner JS. 2015. Experimental inoculation of Egyptian rousette bats (*Rousettus aegyptiacus*) with viruses of the Ebolavirus and Marburgvirus genera. Viruses 7:3420–3442.

11. Baker ML, Schountz T, Wang LF. 2013. Antiviral immune responses of bats: A review. Zoonoses Public Hlth 60:1–13.

12. Zhang G, Cowled C, Shi Z, Huang Z, Bishop-Lilly KA, Fang X, Wynne JW, Xiong Z, Baker ML, Zhao W, Tachedjian M, Zhu Y, Zhou P, Jiang X, Ng J, Yang L, Wu L, Xiao J, Feng Y, Chen Y, Sun X, Zhang Y, Marsh GA, Crameri G, Broder CC, Frey KG, Wang L-F, Wang J. 2013. Comparative analysis of bat genomes provides insight into the evolution of flight and immunity. Science 339:456–460.

13. Zhou P, Tachedjian M, Wynne JW, Boyd V, Cui J, Smith I, Cowled C, Ng JH, Mok L, Michalski WP. 2016. Contraction of the type I IFN locus and unusual constitutive expression of IFN-α in bats. Proc Natl Acad Sci USA:201518240.

14. Hayward JA, Tachedjian M, Cui J, Cheng AZ, Johnson A, Baker ML, Harris RS, Wang L-F, Tachedjian G. 2018. Differential evolution of antiretroviral restriction factors in pteropid bats as revealed by *APOBEC3* gene complexity. Mol Biol Evol 35:1626–1637.

15. Pavlovich SS, Lovett SP, Koroleva G, Guito JC, Arnold CE, Nagle ER, Kulcsar K, Lee A, Thibaud-Nissen F, Hume AJ. 2018. The Egyptian rousette genome reveals unexpected features of bat antiviral immunity. Cell 173:1098–1110. e18.

16. Mandl JN, Schneider C, Schneider DS, Baker ML. 2018. Going to bat(s) for studies of disease tolerance. Front Immunol 9:2112.

17. Hawkins JA, Kaczmarek ME, Müller MA, Drosten C, Press WH, Sawyer SL. 2019. A metaanalysis of bat phylogenetics and positive selection based on genomes and transcriptomes from 18 species. Proc Natl Acad Sci U S A 116:11351–11360.

18. Hölzer M, Schoen A, Wulle J, Müller MA, Drosten C, Marz M, Weber F. 2019. Virus-and interferon alpha-induced transcriptomes of cells from the microbat *Myotis daubentonii*. iScience 19:647–661.

19. Morrison JH, Miller C, Bankers L, Crameri G, Wang L-F, Poeschla EM. 2020. A potent postentry restriction to primate lentiviruses in a yinpterochiropteran bat. Mbio 11.

20. Jebb D, Huang Z, Pippel M, Hughes GM, Lavrichenko K, Devanna P, Winkler S, Jermiin LS, Skirmuntt EC, Katzourakis A, Burkitt-Gray L, Ray DA, Sullivan KAM, Roscito JG, Kirilenko BM, Dávalos LM, Corthals AP, Power ML, Jones G, Ransome RD, Dechmann DKN, Locatelli AG, Puechmaille SJ, Fedrigo O, Jarvis ED, Hiller M, Vernes SC, Myers EW, Teeling EC. 2020. Six reference-quality genomes reveal evolution of bat adaptations. Nature 583:578–584.

21. Neil SJD, Zang T, Bieniasz PD. 2008. Tetherin inhibits retrovirus release and is antagonized by HIV-1 Vpu. Nature 451:425–430.

22. Sakuma T, Noda T, Urata S, Kawaoka Y, Yasuda J. 2009. Inhibition of Lassa and Marburg virus production by tetherin. J Virol 83:2382–2385.

23. Hoffmann M, Nehlmeier I, Brinkmann C, Krähling V, Behner L, Moldenhauer A-S, Krüger N, Nehls J, Schindler M, Hoenen T. 2019. Tetherin inhibits Nipah virus but not Ebola virus replication in fruit bat cells. J Virol 93.

24. Wynne JW, Wang L-F. 2013. Bats and viruses: Friend or foe? PLoS Pathog 9:e1003651.

25. Wang SM, Huang KJ, Wang CT. 2019. Severe acute respiratory syndrome coronavirus spike protein counteracts BST2-mediated restriction of virus-like particle release. J Med Virol 91:1743–1750.

26. Wang L-F, Anderson DE. 2019. Viruses in bats and potential spillover to animals and humans. Curr Opin Virol 34:79–89.

27. Hayward JA, Tachedjian M, Kohl C, Johnson A, Dearnley M, Jesaveluk B, Langer C, Solymosi P, Hille G, Nitsche A, Sánchez CA, Werner A, Kontos D, Crameri G, Marsh GA, Baker ML, Poumbourios P, Drummer HE, Holmes EC, Wang L-F, Smith I, Tachedjian G. 2020. Infectious KoRV-related retroviruses circulating in Australian bats. Proc Natl Acad Sci U S A 117:9529–9536.

28. Perez-Caballero D, Zang T, Ebrahimi A, McNatt MW, Gregory DA, Johnson MC, Bieniasz PD. 2009. Tetherin inhibits HIV-1 release by directly tethering virions to cells. Cell 139:499–511.

29. Blanco-Melo D, Venkatesh S, Bieniasz PD. 2016. Origins and evolution of tetherin, an orphan antiviral gene. Cell Host Microbe 20:189–201.

30. Cocka LJ, Bates P. 2012. Identification of alternatively translated tetherin isoforms with differing antiviral and signaling activities. PLoS Pathog 8:e1002931.

31. Galão Rui P, Le Tortorec A, Pickering S, Kueck T, Neil Stuart JD. 2012. Innate sensing of HIV-1 assembly by Tetherin induces NFκB-dependent proinflammatory responses. Cell Host Microbe 12:633–644.

32. Tokarev A, Suarez M, Kwan W, Fitzpatrick K, Singh R, Guatelli J. 2013. Stimulation of NF-κB activity by the HIV restriction factor BST2. J Virol 87:2046–2057.

33. Matsuda A, Suzuki Y, Honda G, Muramatsu S, Matsuzaki O, Nagano Y, Doi T, Shimotohno K, Harada T, Nishida E. 2003. Large-scale identification and characterization of human genes that activate NF-κB and MAPK signaling pathways. Oncogene 22:3307–3318.

34. Ishikawa J, Kaisho T, Tomizawa H, Lee BO, Kobune Y, Inazawa J, Oritani K, Itoh M, Ochi T, Ishihara K, Hirano T. 1995. Molecular cloning and chromosomal mapping of a bone marrow stromal cell surface gene, BST2, that may be involved in pre-B-cell growth. Genomics 26:527–534.

35. Liberatore RA, Bieniasz PD. 2011. Tetherin is a key effector of the antiretroviral activity of type I interferon in vitro and in vivo. Proc Natl Acad Sci U S A 108:18097–18101.

36. Erikson E, Adam T, Schmidt S, Lehmann-Koch J, Over B, Goffinet C, Harter C, Bekeredjian-Ding I, Sertel S, Lasitschka F. 2011. In vivo expression profile of the antiviral restriction factor and tumor-targeting antigen CD317/BST-2/HM1. 24/tetherin in humans. Proc Natl Acad Sci U S A 108:13688–13693.

37. Yang H, Wang J, Jia X, McNatt MW, Zang T, Pan B, Meng W, Wang H-W, Bieniasz PD, Xiong Y. 2010. Structural insight into the mechanisms of enveloped virus tethering by tetherin. Proc Natl Acad Sci U S A 107:18428–18432.

38. Hinz A, Miguet N, Natrajan G, Usami Y, Yamanaka H, Renesto P, Hartlieb B, McCarthy AA, Simorre J-P, Göttlinger H. 2010. Structural basis of HIV-1 tethering to membranes by the BST-2/tetherin ectodomain. Cell Host Microbe 7:314–323.

39. Kupzig S, Korolchuk V, Rollason R, Sugden A, Wilde A, Banting G. 2003. Bst-2/HM1. 24 is a raft-associated apical membrane protein with an unusual topology. Traffic 4:694–709.

40. Rollason R, Korolchuk V, Hamilton C, Schu P, Banting G. 2007. Clathrin-mediated endocytosis of a lipid-raft-associated protein is mediated through a dual tyrosine motif. J Cell Sci 120:3850–3858.

41. Takeda E, Nakagawa S, Nakaya Y, Tanaka A, Miyazawa T, Yasuda J. 2012. Identification and functional analysis of three isoforms of bovine BST-2. PLoS One 7:e41483.

42. Nehls J, Businger R, Hoffmann M, Brinkmann C, Fehrenbacher B, Schaller M, Maurer B, Schönfeld C, Kramer D, Hailfinger S. 2019. Release of immunomodulatory Ebola virus glycoprotein-containing microvesicles is suppressed by tetherin in a species-specific manner. Cell Reports 26:1841–1853. e6.

43. Iwabu Y, Fujita H, Kinomoto M, Kaneko K, Ishizaka Y, Tanaka Y, Sata T, Tokunaga K. 2009. HIV-1 accessory protein Vpu internalizes cell-surface BST-2/tetherin through transmembrane interactions leading to lysosomes. J Biol Chem 284:35060–35072.

44. Ruedi M, Stadelmann B, Gager Y, Douzery EJ, Francis CM, Lin L-K, Guillén-Servent A, Cibois A. 2013. Molecular phylogenetic reconstructions identify East Asia as the cradle for the evolution of the cosmopolitan genus *Myotis* (Mammalia, Chiroptera). Mol Phylogenet Evol 69:437–449.

45. Fukuma A, Abe M, Morikawa Y, Miyazawa T, Yasuda J. 2011. Cloning and characterization of the antiviral activity of feline Tetherin/BST-2. PLoS One 6:e18247.

46. Benton MJ, Donoghue PCJ. 2007. Paleontological evidence to date the tree of life. Mol Biol Evol 24:26–53.

47. Periasamy P, Hutchinson PE, Chen J, Bonne I, Shahul Hameed SS, Selvam P, Hey YY, Fink K, Irving AT, Dutertre C-A, Baker M, Crameri G, Wang L-F, Alonso S. 2019. Studies on B cells in the fruit-eating black flying fox (*Pteropus alecto*). Front Immunol 10.

48. Mok L, Wynne JW, Ford K, Shiell B, Bacic A, Michalski WP. 2015. Proteomic analysis of *Pteropus alecto* kidney cells in response to the viral mimic, Poly I:C. Proteome Sci 13:1–11.

49. Doering TL, Englund PT, Hart GW. 2001. Detection of glycophospholipid anchors on proteins. Curr Protoc Protein Sci:12.5. 1–12.5. 14.

50. Li F, Li C, Wang M, Webb GI, Zhang Y, Whisstock JC, Song J. 2015. GlycoMine: a machine learning-based approach for predicting N-, C- and O-linked glycosylation in the human proteome. Bioinformatics 31:1411–1419.

51. Dubé M, Roy BB, Guiot-Guillain P, Mercier J, Binette J, Leung G, Cohen ÉA. 2009. Suppression of Tetherin-restricting activity upon human immunodeficiency virus type 1 particle release correlates with localization of Vpu in the trans-Golgi network. J Virol 83:4574–4590.

52. Dietrich I, McMonagle EL, Petit SJ, Vijayakrishnan S, Logan N, Chan CN, Towers GJ, Hosie MJ, Willett BJ. 2011. Feline tetherin efficiently restricts release of feline immunodeficiency virus but not spreading of infection. J Virol 85:5840–5852.

53. Arnaud F, Black SG, Murphy L, Griffiths DJ, Neil SJ, Spencer TE, Palmarini M. 2010. Interplay between ovine bone marrow stromal cell antigen 2/tetherin and endogenous retroviruses. J Virol 84:4415–4425.

54. Wang LF, Anderson DE. 2019. Viruses in bats and potential spillover to animals and humans. Curr Opin Virol 34:79–89.

55. Teeling EC, Vernes SC, Dávalos LM, Ray DA, Gilbert MTP, Myers E, Consortium BK. 2018. Bat biology, genomes, and the Bat1K project: to generate chromosome-level genomes for all living bat species. Annu Rev Anim Biosci 6:23–46.

56. O’Shea TJ, Cryan PM, Cunningham AA, Fooks AR, Hayman DTS, Luis AD, Peel AJ, Plowright RK, Wood JLN. 2014. Bat flight and zoonotic viruses. Emerg Infect Dis 20:741–745.

57. Kacprzyk J, Hughes GM, Palsson-McDermott EM, Quinn SR, Puechmaille SJ, O’Neill LA, Teeling EC. 2017. A potent anti-inflammatory response in bat macrophages may be linked to extended longevity and viral tolerance. Acta Chiropt 19:219–228.

58. Celestino M, Calistri A, Del Vecchio C, Salata C, Chiuppesi F, Pistello M, Borsetti A, Palù G, Parolin C. 2012. Feline tetherin is characterized by a short N-terminal region and is counteracted by the feline immunodeficiency virus envelope glycoprotein. J Virol 86:6688–6700.

59. Schubert HL, Zhai Q, Sandrin V, Eckert DM, Garcia-Maya M, Saul L, Sundquist WI, Steiner RA, Hill CP. 2010. Structural and functional studies on the extracellular domain of BST2/tetherin in reduced and oxidized conformations. Proc Natl Acad Sci U S A 107:17951–17956.

60. Andrew AJ, Miyagi E, Kao S, Strebel K. 2009. The formation of cysteine-linked dimers of BST-2/tetherin is important for inhibition of HIV-1 virus release but not for sensitivity to Vpu. Retrovirology 6:80.

61. McNatt MW, Zang T, Hatziioannou T, Bartlett M, Fofana IB, Johnson WE, Neil SJ, Bieniasz PD. 2009. Species-specific activity of HIV-1 Vpu and positive selection of tetherin transmembrane domain variants. PLoS Pathog 5:e1000300.

62. Lim ES, Malik HS, Emerman M. 2010. Ancient adaptive evolution of tetherin shaped the functions of Vpu and Nef in human immunodeficiency virus and primate lentiviruses. J Virol 84:7124–7134.

63. Liu J, Chen K, Wang J-H, Zhang C. 2010. Molecular Evolution of the Primate Antiviral Restriction Factor Tetherin. PLoS One 5:e11904.

64. Venkatesh S, Bieniasz PD. 2013. Mechanism of HIV-1 virion entrapment by tetherin. PLoS Pathog 9:e1003483.

65. Andrew AJ, Kao S, Strebel K. 2011. C-terminal hydrophobic region in human bone marrow stromal cell antigen 2 (BST-2)/tetherin protein functions as second transmembrane motif. J Biol Chem 286:39967–39981.

66. Krogh A, Larsson B, Von Heijne G, Sonnhammer EL. 2001. Predicting transmembrane protein topology with a hidden Markov model: application to complete genomes. J Mol Biol 305:567–580.

67. Sakuma T, Sakurai A, Yasuda J. 2009. Dimerization of tetherin is not essential for its antiviral activity against Lassa and Marburg viruses. PLoS One 4:e6934.

68. Hayward JA, Tachedjian M, Cui J, Field H, Holmes E, Wang L-F, Tachedjian G. 2013. Identification of diverse full-length endogenous betaretroviruses in megabats and microbats. Retrovirology 10:35.

69. Skirmuntt EC, Escalera-Zamudio M, Teeling EC, Smith A, Katzourakis A. 2020. The potential role of endogenous viral elements in the evolution of bats as reservoirs for zoonotic viruses. Annu Rev Virol 7:103–119.

70. Cui J, Tachedjian G, Tachedjian M, Holmes EC, Zhang S, Wang L-F. 2012. Identification of diverse groups of endogenous gammaretroviruses in mega and microbats. J Gen Virol 93:2037–2045.

71. Hause BM, Nelson EA, Christopher-Hennings J. 2020. North American big brown bats (Eptesicus fuscus) harbor an exogenous deltaretrovirus. mSphere 5.

72. Murphy L, Varela M, Desloire S, Ftaich N, Murgia C, Golder M, Neil S, Spencer TE, Wootton SK, Lavillette D. 2015. The sheep tetherin paralog oBST2B blocks envelope glycoprotein incorporation into nascent retroviral virions. J Virol 89:535–544.

73. Swiecki M, Scheaffer SM, Allaire M, Fremont DH, Colonna M, Brett TJ. 2011. Structural and biophysical analysis of BST-2/tetherin ectodomains reveals an evolutionary conserved design to inhibit virus release. J Biol Chem 286:2987–2997.

74. Yang Z. 2007. PAML 4: phylogenetic analysis by maximum likelihood. Mol Biol Evol 24:1586–1591.

75. Kumar S, Stecher G, Suleski M, Hedges SB. 2017. TimeTree: A resource for timelines, timetrees, and divergence times. Mol Biol Evol 34:1812–1819.

76. Pierleoni A, Martelli P, Casadio R. 2008. PredGPI: a GPI-anchor predictor. BMC Bioinformatics 9:1.

77. Edgar RC. 2004. MUSCLE: multiple sequence alignment with high accuracy and high throughput. Nuc Acid Res 32:1792–1797.

78. Gasteiger E, Hoogland C, Gattiker A, Wilkins MR, Appel RD, Bairoch A. 2005. Protein identification and analysis tools on the ExPASy server. The proteomics protocols handbook:571–607.

79. Meri S, Lehto T, Sutton CW, Tyynelä J, Baumann M. 1996. Structural composition and functional characterization of soluble CD59: heterogeneity of the oligosaccharide and glycophosphoinositol (GPI) anchor revealed by laser-desorption mass spectrometric analysis. Biochem J 316:923–935.

80. Cowled C, Baker M, Tachedjian M, Zhou P, Bulach D, Wang L-F. 2011. Molecular characterisation of Toll-like receptors in the black flying fox *Pteropus alecto*. Dev Comp Immunol 35:7–18.

81. Sambrook J, Fritsch EF, Maniatis T. 1989. Molecular cloning, vol 2. Cold spring harbor laboratory press New York.

82. Brown TA. 2002. Synthesis and processing of RNA, Genomes. Oxford: Wiley-Liss.

83. Kent WJ. 2002. BLAT—the BLAST-like alignment tool. Genome Res 12:656–664.

84. Livak KJ, Schmittgen TD. 2001. Analysis of relative gene expression data using real-time quantitative PCR and the 2− ΔΔCT method. Methods 25:402–408.

85. Irving AT, Ng JHJ, Boyd V, Dutertre C-A, Ginhoux F, Dekkers MH, Meers J, Field HE, Crameri G, Wang L-F. 2020. Optimizing dissection, sample collection and cell isolation protocols for frugivorous bats. Methods Ecol Evol 11:150–158.

86. Andrews S. 2010. FastQC: a quality control tool for high throughput sequence data. http://www.bioinformatics.babraham.ac.uk/projects/fastqc/. Accessed 25/07/2022.

87. Jouvenet N, Neil SJD, Zhadina M, Zang T, Kratovac Z, Lee Y, McNatt M, Hatziioannou T, Bieniasz PD. 2009. Broad-spectrum inhibition of retroviral and filoviral particle release by tetherin. J Virol 83:1837–1844.

