## Supplementary material for "Unique Evolution of Antiviral Tetherin in Bats": SI Data 1

[a]Health Security Program, Life Sciences Discipline, Burnet Institute, Life Sciences, Melbourne, VIC 3004, Australia; [b]Department of Microbiology, Monash University, Clayton, VIC 3168, Australia; [c]CSIRO, Australian Centre for Disease Preparedness, Health and Biosecurity Business Unit, Geelong, VIC 3220, Australia; [d]Zhejiang University-University of Edinburgh Institute, Zhejiang University School of Medicine, Zhejiang University International Campus, China; [e]Second Affiliated Hospital, Zhejiang University School of Medicine, Hangzhou, China; [f]Programme in Emerging Infectious Diseases, Duke-NUS Medical School, Singapore, Singapore; [g]CAS Key Laboratory of Molecular Virology & Immunology, Institut Pasteur of Shanghai, Chinese Academy of Sciences, China; [h]Center for Biosafety Mega-Science, Chinese Academy of Sciences, China; [i]Singhealth Duke-NUS Global Health Institute, Singapore; [j]Department of Microbiology and Immunology, The University of Melbourne, at The Peter Doherty Institute for Infection and Immunity, Melbourne, VIC 3000, Australia.

**This PDF file includes:**

SI Data 1

### SI Data

**Data SI 1**. Multiple sequence alignment of 27 bat tetherin protein homologues of human l-tetherin.

```
>Tetherin_(Cynopterus_sphinx)
MAPTWYHYFPVPMGDYSEKPALRDRKLP---GWLWIPLVLV---GLGLLV
A---LIIFIIKANSEACKDGLRAEQKCRNETHLLELQLTRAQESLVAAKA
QAASCNQTVETLKSSLEMEKAESQKQQKLAQELQGEIRNLKQELEN----
------------------------TAA-------ELEQLRKEHEKN----
-GEKNGSTSF------RDARSSLVVAVLLGLSFGALLA
>Tetherin_(Pteropus_alecto)_Isoform_A
MAPTLYHYFPVPMDDHSKKVVLGNRNLP---RWLWILLVLV---ILGLTV
A---VIVLAVENSSEACKNGLQAEQKCRNETHLLKLQLTQTQESLGVAKA
QAASCNQTVGTLNSSLEMEKAESQKQRELAQKLQGEITNLTQQLKN----
------------------------TAA-------ELEQLRKEHAS-----
-GEKNGSTSS------RNARSSLVVVVLLSLSFRALLA
>Tetherin_(Pteropus_vampyrus)
MAPTLYHYFPVPMDDHSKKVVLGDRKLP---GWLGILLVLV---ILGLIV
A---VIVLAVKANSEACKDGLQAEQKCRNKTHLLELQLTQAQESLGVAKA
QAASCNQTVGTLKSSLEMEKAESQKQRELAQELQGEITNLTQQLKN----
------------------------TAA-------ELEQLRKEHAS-----
-GEKNGSTSS------RNAGSSLVVVVLLSLSFRALLA
>Tetherin_(Eidolon_helvum)
MAPTWYHYFPVPMDDHSEKLKLGNRKLP---GWLCILLVLV---ALGLLV
P---LIVFIIKANSEACKDGLRAEEKCRNKTDLLELQLTQAQESLVVAKD
QTALCNQTVGTLKLSLEMEKAESQKQRELAQELQGEIRNLKQKLEN----
------------------------TAA-------ELEQLRKEHAS-----
-GEKNGATSS------RNARSSLVVAVLLSLSFIALLA
>Tetherin_(Eonycteris_spelaea)
MAPTLYHYFPVPMDDHPKKRVLGNRKLP---KWLQILLVLA--ALLSLLV
P---LIFFAIKANSEACKDGLLAEQKCQNKTHLLELQLTQAQESLVVAED
QAVLCNRTVATLNSSLEMEKAESQKQRELAQELQGEIWNLKQKLEN----
------------------------TMA-------ELEQLRKEHES-----
-DEKNGSTSS------GNARSSLVVAVLLSLSFGALLA
>Tetherin_(Rousettus_aegyptiacus)
MAPALYHYFPVPMDDHSEKVVLRNRKML---KWLWILPVLA--LVLSLLV
A---LIVFAIEANSKACKDGLLAEQKCLNKTRLLELQLTQAQESLVVAEA
QAYLCNQTVGTLKSSLEMEKDESQKQRELAQKLQGENGNLKQELEN----
------------------------MMA-------ELEQLRKEHAS-----
-DEKNGSTSS------RNARSFLVVAVLLSLSFGALLA
>Tetherin_(Myotis_lucifugus)
MAPTFYCHPPVLMDEYPKK--MGDRKLPVREG---ILLGLL---VVGLFV
A---VGVLAAKANSPACKDGLRAEQECRNNTHLLEQELTQAQEVLRGTEA
QAATCNQTVETLKISLKEEKAAGHKQQELVQKLQEEIKTLNQSLQN----
------------------------KSSELEKKSAELEQLRKENEAL----
-VSAKGPPDS------GVSLHLSMAAVLLPLSLLVLLA
>Tetherin_(Myotis_brandtii)
MAPTFYCHPTVPMDEYPKK--LGDRKLPVREGIVGILLVVV----VGLSV
A---VGVLAAKANSPACKDGLPAEQECRNITHLLEQELTQAQEVLRETEA
QAATCNQTVETLKISLKEEKAAGHKQQELVQKLQEEIKTLNQSLQD----
------------------------KSAELEEKSSELEQLRKENEDL----
-VSAKGPPNS------GITLRLSVAAVLLSLSLLTLLA
>Tetherin_(Myotis_davidii)
MAPTSYCHPPVPMDEYPKK--LGDRKLPVREGIVGILLGLL---VVGLSV
A---VGVLAAKANSPACNDGLRAEQECRNITHLLEQELTQAQEVLRGTEA
QSATCNQTVETLKISLKEEKAVGLRQQELVQKLQEEIKTLNQSLKD----
```

```
------------------------KSAELEEKSTELGKLRKENEAL----
-VSAKGPPNS------GITLSFSVVAVLLPMSLLALLA
>Tetherin_(Myotis_myotis)
MAPTFYCHPPVPMDEYPKK--LGERKLPVREGIVGILLGLL---VVGLSV
A---VGVLAAKANSPACKDGLQAEQECRNITHLLEQELTQAQEVLRGTEA
QAATCNQTVETLKISLKEEKAAGHKQQELVQKLQEEIKTLNQSLRD----
------------------------KSADLEKKSAELEQLRKENEAL----
-VSAKGPPNS------GITLSLSVAAVLLPLSLQALLA
>Tetherin_(Myotis_laniger)
MAPTFYCHPPVPMDEYPKK--LGDRKLPVREGIVWILLGLL---VVGLSV
A---VGVLAAKANSPACKDGLLAEQECRNITHLLEQELTQAQEVLRGTEA
QAATCNQTVETLKISLKEEKAAGHRQQELVQKLQEEIKTLNQSLQE----
------------------------KSAELEKKSEELEQLRKENEAL----
-VSAKGPPNS------GITLSLSVVAVLLPMSLLALLA
>Tetherin_(Myotis_macropus)
MAPTFYCHPPVPMDEYPKK--LGDQKLPVREGIVWILLGLL---VVGLSV
A---VGVLAAKANSPACKDGLLAEQECRNITHLLEQERTQAQEVLRGTEA
QAATCNQTVETLKISLKEEKAAGHRQQELVQKLQEEIKTLNQSLQE----
------------------------KSAELEKKSEELEQLRKENEAL----
-VSAKGPPNS------GITLSLSVVAVLLPMSLLALLA
>Tetherin_(Myotis_ricketti)
MAPTFYCHPPVPMDEYPKK--LGDQKLPVREGIVWILLGLL---VVGLSV
A---VGVLAAKANSPACKDGLLAEQECRNITHLLEQERTQAQEVLRGTEA
QAATCNQTVETLKISLKEEKAAGHRQQELVQKLQEEIKTLNQSLQE----
------------------------KSAELEKKSEELEQLRKENEAL----
-VSAKGPPNS------GITLSLSVVAVLLPMSLLALLA
>Tetherin_(Murina_leucogaster)
MAPTFYCHPPVPMDEYPKK--LGDRKLPVREGIVGILLGLL---VVSLSV
A---VGVLAAKANSPACKDGLRAEQECRNITHLLEQELTQAQEVLRGTEA
QAATCNQTVETLKISLKEEKAAGHRQQELVQKLQEEIKTLNQELQE----
------------------------KSA-------ELEQLRKENEAL----
-VSAKGPPNS------GITLSLSVVAVLLPMSLLALLA
>Tetherin_(Hipposideros_armiger)
MAPISYKYCPVSMDDNLSP--LQSRKLP---WWLWILLAVVLMLVLILIV
L---TIFFAVRANSKACKDGLQAEQECRNFTHLLEHQLTQANKVLLDAEI
WAATCNKTVETLTASLEMEKARGHKQQELVQKLQGEIAELKQKLQD----
------------------------TSK-------ELDRLRKETETS----
-DTGNGSTTGNGSTSFGDTINLFVVSVLLTLSLGALLA
>Tetherin_(Rhinolophus_ferrumequinum)
MAPISCHYRPVPMGDKLNS-VLGGRKAP---CWLVIMLVVG---VGILLV
P---LSIFAVKANRESCKDGLLAEQKCQNLTHLLEHQLTQAHNLLLKMKN
QTATCNQTVVTLKASLEAEKAQGPKQQELLQELQGEIKKLKQKLQD----
------------------------TTT-------ELKQLRKKNEAS----
-EEGKGSTSS------GNTINPLVIHVLLTLSLRALLA
>Tetherin_(Rhinolopus_macrotis)
MAPVSCHYRPVSMDDKLTS-VFGGHRPS---WWMVIVLGVA---VVILFV
V---TSIFAAMANREACKDGLLAEQKCQNFTHLLERQLTQAQNVLLGMKI
QAATCNQTVVTLTASLETEKAHGHKQQELVQQLQGEIEKLKQKLQD----
------------------------TTT-------ELEQLRKKNDAS----
-EGGNGSTSS------GNTINSLAIYVLLTLSLGALLA
>Tetherin_(Taphozous_melanopogon)
------------MDDYLEKLKLWDCKLP---RWLGILLLLL---VVGEFV
L---LIIFGVRDSSNVCKDSLRAEQECRNITHLLQHQLTQAQEVVLETEA
QAASCNKTVVTLMESLEKEKAQGHKQQELVQELQEEIEKLKQELQD----
------------------------TGTEL-----ELERLRKKDESW----
-GLKTNSNS-------GNILRPLAITVLLTLCLRTLLA
>Tetherin_(Carollia_brevicauda)
MTSTFYHYTLLPMGKNSRELMLGYYQLPRQLKILLALLGLL---VVGLSV
```

A---TTIFAVQAFSPACKDGRRAEQECRNFTHLLQRQLTQAQDILLKTEA
QAATYNQTAETLMASLKVEQAQGQKLQEQVQELQEEIETLKQKLQDTTQK
LQDTTQKLQD--------------TTQKLQGTTAELNQLRKDHESC----
---GYDSTNS------GNTLGLSVVAMLLTLSLLDLLA
>Tetherin_(Carollia_perspicillata)
MTSTFYHYTLLPMGKNSRELMLGYYQLPRQLKILLALLGLL---VVGLSV
A---TTIFAVQAFSPACKDGRRAEQECRNFTHLLQRQLTQAQDILLKTEA
QAATYNQTAETLMASLKVEQAQGQKLQEQVQELQEEIETLKQKLQDTTQK
LQDTTQKLQD--------------TTQKLQGTTAELNQLRKDHESC----
---SYDSTNS------GNTLGLSVVAMLLTLSLLDLLA
>Tetherin_(Desmodus_rotundus)
MTSTFYHYIPLPMSENSRELMLEYHKLP---RWLGILLALLVLLVVGLLV
A---TIILAVQAHSPACKDGRRAEQECRNFTHLLESQRTRAQEILLKTKA
QAATYNQTVVTLMASLKVEQAQGQKLREQVQELQEEIKTLKQKLQDTTQK
LQVTTTELQNTTQKLQVTTAKLQVTTQKLQDTTTELNQLRKDHESF----
-GWGNGSTSS------GNTLSLSVVVLLLTLSLLDLLA
>Tetherin_(Macrotus_californicus)
MAPTFYHYVPLPMGDNSSKLMLRGYKLP---KWLVILLALLVLLVVGLLL
A---TITFAVQAHRPACKDGHRAEQECRNFTHLLESQQTQTQEILLKTKA
EAATYNQTVVTLMASLKVEQAQGQKLQEQVQELQEEIKTLKQKLQD----
------------------------TTQKLQDTTTELNQLREDQESF----
-SRGHGSTSS------GSTLSLSVVTMLLTLRLLDLLA
>Tetherin_(Artibeus_jamaicensis)
MTSTFYHYTSLPSDENSREQMQGHYKIP---LWLSILLALL---GTGLVV
AMALAIIFYNQAHSLACMDGFKMEQECQNVTNSLKSKLAQTQEILLKTEA
QAATYNQTAVTLMASLKVEQAQGQMLQKQVQELQKEIKTLKQNLQD----
------------------------TTQKLQDTTQKLNQLRKDHESS----
-GPGKGPTDS------GNTLSLSMISMLLTLSLLDLLA
>Tetherin_(Uroderma_bilobatum)
MTSTFYHYISLPLDENSREQMQGHYKMP---RWLLILLALL---VVGLVV
ASALAIYFAHQAHSPACKNGLKVEQECQNVTDSLKSKLAQTQEILLKTEA
QAATYNQTVAAQMASLKAERAQSQKLQEQVQELQKEIKTLKQKLQD----
------------------------TTQKLQDTTAELNQLRKDHGSS----
-SRGNGPTSS------ENTLSLSVVAMLLTLSLLDLLA
>Tetherin_(Tadarida_basiliensis)
MAPTFY-YRPLSMDECSEYGKQQYWRLP---RCLVITVFLM---LVSMSV
T---VIVLAVKLNSTACKNGLQAEQECRNRTHLLENQLTQTQ---LKTEV
QAATCNQTVATLMASLKKEKAQSDTQQELVQTLQEKIKELKQKLNT----
------------------------SPD---------QGLRKENEAS----
-GGENSSTSF------GNTLSRCVVTLLLAVSLQVLLA
>Tetherin_(Miniopterus_natalensis)
MAGTLYHHSPVPMDDRSNNLMLWCHRWL---KWLLVIPVMV-----VLIV
L---VIILANRLNSITCKDVLGAEQECQNITY-LEHQLNQAQRVLLETN-
--ATCNQTVVTLMDSLKMEQAQGHKQQQLVQKLQEEIKELKQNVQN----
------------------------TSM-------ELERLRKWKETS----
-GREEDSTSS------GNTFSLCMVIVLLPLTLEVLLA
>Tetherin_(Miniopterus_schreibersii)
MAGTLYHHSPVPMGDRSNNLMLWCHRWL---KWLLVIPVMV-----VLIV
L---VIILANRPNGITCKDVLRAEQECQNITY-LEHQLNQAQRVLLETN-
--ATCNQTVVTLMDSLKMEQAQGHKQQQLVQKLQEEIKELKQNLQN----
------------------------TSM-------ELERLRKWKETS----
-GREEDSTSS------GNTFSLCMVIVLLPLTLEVLLT
